# Development of Pial Collaterals by Extension of Pre-existing Artery Tips

**DOI:** 10.1101/2023.12.07.570539

**Authors:** Suraj Kumar, Swarnadip Ghosh, Niloufer Shanavas, Vinayak Sivaramakrishnan, Manish Dwari, Soumyashree Das

## Abstract

Pial collaterals provide protection from ischemic damage and improve prognosis of stroke patients. The origin or precise sequence of events underlying pial collateral development is unclear and has prevented clinicians from adapting new vascularization and regeneration therapies. We use genetic lineage tracing and intra-vital imaging of mouse brains at cellular resolution to show that during embryogenesis, pial collateral arteries develop from extension and anastomoses of pre-existing artery tips, in a VegfR2 dependent manner. This process of artery tip extension occurs on pre-determined microvascular tracks. Our data demonstrate that an arterial receptor, Cxcr4, earlier shown to drive artery cell migration and coronary collateral development, is dispensable for formation and maintenance of pial collateral arteries. Our study shows that collateral arteries of the brain are built by a unique mechanism, distinct from that of the heart.

## INTRODUCTION

Pial (leptomeningeal) collateral arteries are an extensive network of inter-artery anastomoses found in the pial layer of the brain. In humans, the extent of pial collateral circulation is a predictor of stroke outcome.^1–3^ Stroke occurs when blood supply is obstructed to a part of the brain and is one of the leading causes of morbidity worldwide (World Health Organization).^4^ Pial collaterals serve as an alternate route of blood flow and alleviate the symptoms of ischemic stroke by bypassing vascular occlusions. Thus, understanding mechanisms which drive the origin and remodelling of pial collateral arteries is of immense clinical relevance.

Results from preclinical studies indicate wide-spread variation in the pial collateral artery network among individuals and organs.^5^ These variations are a result of differential expression and function of proteins associated with Vascular endothelial growth factor (Vegf) signalling axis.^6–8^ Different mouse strains demonstrate differential Vegfa expression level which corelate with their pial collateral density.^9^ Similarly, global overexpression of Vegf ligand increases number of native pial collaterals and improve collateral perfusion post-injury.^6,7^ The receptor mediating the function of Vegf ligand during the formation of pial collaterals is Vegf receptor-2 (VegfR2).^7^ A global knockdown of Vegf or VegfR2 during embryogenesis and in a time-frame when pial collaterals are expected to form, results in reduction in their number and diameter.^7^ Thus, Vegf is a key driver of pial collateral formation and a critical factor which regulates variation in extent of pial vascular network in mice.

The vital role of Vegf in murine pial collateral development was discovered almost a decade ago.^6,7^ However, it is still unclear how the various molecular players of Vegf pathway orchestrate the formation or remodelling of pial collateral network. In mice, it is hypothesized that an *arteriole-specific sprouting angiogenesis like* process leads to the formation of the foremost pial collaterals.^7^ However, molecular evidence supporting the identity of these sprouting cells (if arteries or capillaries) is missing. Additionally, the cellular origin or fate of these sprouts is unknown. Finally, the possibility of cooperation from other vascular processes, such as arterialization, arteriogenesis or artery-reassembly, in building pial collaterals, needs to be thoroughly investigated.

In this study, we use genetic lineage tracing in conjunction with live imaging of pial vasculature of transgenic mouse embryos and adults to understand mechanisms by which Vegf regulates pial collateral development and maintenance. Our data indicate that the cellular source of pial collateral arteries is primarily pre-existing artery endothelial cells (ECs) which build the first network of pial collaterals at embryonic day (e)15.5 via VegfR2-dependent artery tip extension. The artery tips extend on pre-determined microvascular tracks which then connect and build pial collaterals. While arterial VegfR2 was essential for formation of pial collaterals *de novo*, it was dispensable for its pruning or maintenance through adulthood. Arterial Cxcr4 (C-X-C chemokine receptor type-4), which is critical for coronary artery formation in mice^10^ was dispensable during pial collateral formation and its maintenance. Together, our data indicate that artery endothelial cells build collateral arteries by utilizing distinct vascular mechanisms in heart and brain.

## RESULTS

### Pial collateral arteries form during embryogenesis

We developed a whole brain imaging technique to reliably visualize all pial arteries (Figures **S1A** and **S1B**). Due to lack of unique markers, accurately identifying pial collateral arteries is exclusively dependent on their location. Pial collateral arteries are artery segments which interconnect the ends of cerebral arteries and can be seen in the watershed region (Figure **S1B**). To acquire this spatial information, it was imperative to image the entire pial artery network. Our technique allowed us to precisely identify pial collateral arteries. We specifically focused on collateral arteries which connected and received blood flow from Anterior Cerebral Artery (ACA) and Middle Cerebral Artery (MCA), as the coverage of this arterial network is most extensive.

We first investigated the developmental stage when pial collaterals first appeared. Mouse brains were harvested at different developmental time points and whole brain immunostaining was performed for Connexin-40 (Cx40) protein, a known marker for mature artery endothelial cell. Analyses of pial arterial network in both hemispheres revealed the appearance of MCA at embryonic day (e)12.5 followed by ACA at e13.5 (Figures **S1C**, **S2A** and **S2B**). At e14.5, both cerebral arteries were present with no arterial connections between them (Figure **S2C**). Subsequently, the first arterial anastomoses appeared at e15.5 (Figure **S2D**) which then remodelled to expand in number through birth/postnatal day (P)0 (Figures **S1C** and **S2D**-**S2F**). This method confirmed that both MCA and ACA are present at e14.5 (Figure **S2G**) and the foremost pial collaterals appear at e15.5 (Figure **S2H**).

To confirm the above results, we next used a knock-in mouse line (*Cx40^eGFP/+^*) where eGFP expression was driven by *Cx40* promoter activity. Brains from different developmental stages confirmed our findings from Cx40 immunostaining. MCA appeared at e12.5, ACA appeared at e14.5 and collateral arteries between MCA and ACA appear at e15.5 (Figures **S3A**-**S3F** and **S1C**). eGFP^+^ cerebral arteries (MCA and ACA) were formed by e14.5 (Figure **S3G**) and the first eGFP^+^ collateral artery between MCA and ACA appeared at e15.5 (Figure **S3H**). Thus, using two independent methods— Cx40 immunostaining and detecting its reporter activity, we demonstrate that mouse pial collateral arteries form during development, and appear precisely at e15.5 (Figure **S1C**).

### Intra-vital imaging reveals arterial origin of pial collaterals by artery tip extension

Despite several hypotheses, there is lack of compelling data in support of vascular mechanisms which drive pial collateral development. Pial collaterals appear at late embryonic stages. At this developmental stage, rapid vascular changes make it difficult to distinguish collateral-specific events from overall vascular growth, thus making it challenging to draw information from fixed brain samples. Additionally, late embryonic stages cannot be cultured *ex vivo* due to extremely demanding metabolic and physiological needs. Thus, we custom-developed an imaging paradigm to perform intra-vital imaging of pial layer at late embryonic stages (e16-e17) (Figure **1A**), a developmental stage where the brain is actively building pial collaterals. Briefly, embryos of the desired developmental time points were dissected out (with their intact yolk sac) from the anesthetized pregnant dams while still connected to the mother via their umbilical cord. Embryos were maintained at 37°C and immobilized at appropriate orientation to aid imaging. The embryos were subjected to stereoscopic or confocal imaging for several hours. This imaging set up allowed visualization of the entire hemisphere─ MCA, ACA, and connected collateral arteries (Figure **1B**).

**Figure 1:**
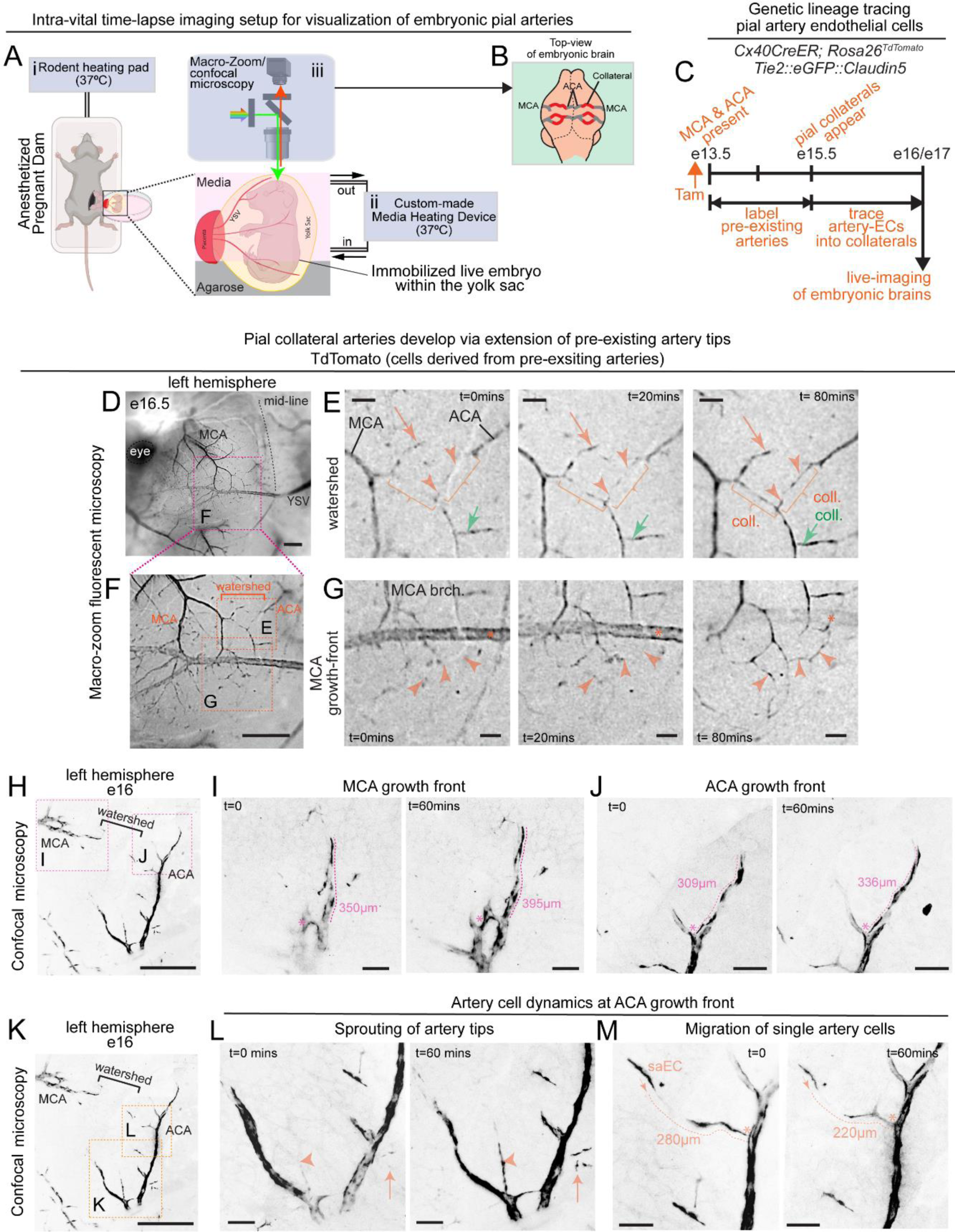
Live imaging of embryonic pial vasculature. (**A**) Intra-vital imaging set up to visualize the developing pial vasculature in mouse embryos. **i** heating pad to maintain body temperature of anaesthetized pregnant dam at 37°C. **ii** custom-built media heating device which also circulates dissection media at 37°C **iii** shows macro-zoom or confocal microscopy performed on live, immobilized embryos. (**B**) Representative schematic of an embryonic brain. Grey and Red show cerebral arteries and pial collaterals respectively. (**C**) Experimental design to genetically label and trace the dynamics of pre-existing artery endothelial cells in a developing pial layer. (**D**-**G**) Images obtained through macro-zoom fluorescent microscopy of e16.5 embryos. In black are cells derived from pre-existing artery endothelial cells. Images were taken by focusing in and out to capture the entire curvature of the embryonic brain. **D** shows an entire left hemisphere. **E** represents watershed region spanned by MCA and ACA artery tips and imaged at time=0 minutes, 20 minutes and 80 minutes respectively. Arrows and arrowheads show growing ends of pre-existing artery tips from MCA and ACA. Brackets mark regions where collaterals appear. **F** shows inset in **D**. **E** and **G** show insets in **F**. **G** represents the MCA growth front imaged at time=0 minutes, 20 minutes and 80 minutes respectively. * shows a superficial yolk sac vessel which photobleaches with time. Arrowheads mark extension and anastomoses of MCA artery tips. (**H**-**J**) Images obtained through confocal microscopy from e16 embryos. **H** shows pial watershed and the cerebral arteries spanning the watershed. **I**, **J** show MCA and ACA growth fronts respectively imaged at time=0 minutes and 60 minutes. In black are cells derived from pre-existing artery endothelial cells. Images shown in **I** are maximum intensity projections from 50 optical slices and capture the entire width of the artery. Images shown in **J** are maximum intensity projections from 40 optical slices and capture the entire width of the artery. * represents an internal landmark from where the length measurements were done. (**K**-**M**) Images obtained through confocal microscopy from e16 embryos. (**L**, **M**) Insets from **K** showing artery cell growth of developing ACA in 60 mins. (**L**) Arrowhead points to an artery tip emerging within 60mins. Arrow points to a region where single recombined artery cells appear. (**M)** Arrow shows direction of migration of single recombined artery cell. Dotted line is the length of micro-vessel between the single recombined cell and the pial artery. MCA, Middle cerebral artery; ACA, Anterior cerebral artery; e, embryonic day; Tam, Tamoxifen; EC, endothelial cell; YSV, yolk sac vessel; brch., branch; coll, collateral; MV, microvascular. Scale bars: **D**, **F**, 1mm; **E**, **G**, 200 µm; **H**, **K**, 500 µm; **I**, **J**, **L**, **M**, 100 µm.

Pial collaterals appear only after the primary branches of MCA and ACA have developed (Figure **S1C**). We hypothesized that the branches of these cerebral arteries contribute towards formation of pial collaterals. To test this hypothesis, *Cx40CreER; Rosa26^TdTomato^*; *Tie2::eGFP::Claudin5* mice were used to label artery ECs with TdTomato and all endothelial cells with eGFP. To activate TdTomato, Tamoxifen was administered at e13.5, when both MCA and ACA are present and would be labelled in 2 days, ^10,11^ by e15.5. Imaging of endothelial cells derived from arteries was performed at e16-e17. This experimental strategy allowed visualization of cellular dynamics specific to artery cells derived from pre-existing arteries (Figure **1C**) and excluded the possibility of capturing other vascular processes.

We performed intra-vital time-lapse imaging of pial arteries and investigated if pial collaterals originate from pre-existing artery endothelial cells. We focused on the watershed region: an initial artery-free region which spans the distal ends of both arterial trees (MCA and ACA) and is later occupied by pial collaterals (Figures **S1C** and **1D**-**1F**). Stereoscopic images of e16.5 brains revealed that branches from MCA and ACA extend (arrowheads, Figure **1E**) and anastomose to build collaterals in the pial watershed (brackets, Figure **1E**). Additionally, TdTomato expression in artery tips (orange arrow, Figure **1E**) and collaterals (green arrow, Figure **1E**) increased over time. These changes were executed within a short span of 80 minutes, and could not have been captured in fixed brain samples.

We next investigated artery cell dynamics associated with the growth fronts of the two cerebral arteries (Figures **1F** and **1G**). At e16.5, the distal branches of the developing MCA extended and anastomosed to build an artery network *de novo* (arrowheads, Figure **1G**). This is the first evidence demonstrating that pre-existing cerebral artery tips build pial collaterals. We then performed intra-vital time-lapse confocal imaging of brains at e16, a stage prior to when we observed collateral artery networks being built *de novo* (Figures **1E** and **1G**). Confocal imaging at e16 provided the necessary cellular resolution to quantitatively assess artery-tip extensions from MCA and ACA branches (Figures **1H**-**1J**). As expected, we clearly observed artery tip extension of MCA branch (Figures **1H** and **1I**) and the ACA branch (Figures **1H** and **1J**) within an hour. Additionally, we noted several artery-specific movements within the pial layer. This included the appearance of single recombined artery endothelial cells (arrows, Figures **1K** and **1L**), movement of these single recombined artery cells towards an artery (Figures **1K**, **1M** and **S4A**), and emergence of arterial sprouts (arrowheads, Figures **1K**, **1L and S4B**-**S4D**). Together, we provide direct evidence for the vascular origin of pial collaterals─ pre-existing MCA/ACA artery tips extend and connect with each other to build pial collaterals *de novo*.

### Artery tip extension occurs on pre-determined microvascular tracks

We next investigated if other endothelial cell sub-types participate in, and/or facilitate, pial artery tip extensions during collateral development. To investigate the same, *Cx40CreER; Rosa26^TdTomato^*; *Tie2::eGFP::Claudin5* mouse line was used. *Tie2::eGFP::Claudin5* is a transgenic mouse strain which expresses a fusion protein of eGFP with Claudin5 (a tight junction protein present in all ECs in the central nervous system). This eGFP-Claudin5 protein expression is under the control of *Tie2* promoter/enhancer, making it visible in all endothelial cells.^12,13^ Thus, in *Cx40CreER*; *Rosa26^TdTomato^*; *Tie2::eGFP::Claudin5* mice, TdTomato marked artery ECs while eGFP marked all ECs. Live imaging of pial watershed showed that all artery cell movements, almost always, occurred along microvascular (MV) tracks which appeared structurally more organized/tubular than the surrounding capillaries (arrowheads, Figures **2A**, **2B** and **S5**). These MV tracks were distinct from the surrounding capillary plexus in that they formed direct connections between two arteries, but were not necessarily entirely populated by TdTomato^+^ artery derived endothelial cells (Figure **2B**). Additionally, our data show that these MV tracks originated from Apj^+^ capillary/venous endothelial cells (Figures **S6A**-**S6H**), expressed microvascular marker Endomucin (Figures **S6I**-**S6L**) and lacked arterial marker Cx40 (Figures **S6M**-**S6O**). Together, our data provide support for the non-arterial nature of the MV tracks (Figure **S6P**), and hence, the intermixing of TdTomato^+^ and TdTomato^-^ cells, are not due to uneven Tamoxifen recombination of *Cx40CreER*. We hypothesized that development of MV tracks precedes artery tip extensions and tested two potential mechanisms by which MV tracks could assist artery tip extensions during pial collateral development (Figure **2C**). Firstly, complete formation of MV tracks between two arteries at e16 could be a pre-requisite for initiation of artery tip extension, as often observed at e16. (Figures **2Ci-iv**). Alternatively, MV tracks could appear first, but continuously grow/remodel to allow artery tip extension (Figures **2Ci, v, vi**) and subsequent pial collateral formation (Figure **2Civ**). If the former was true, the number/length of MV tracks in a given watershed would remain unchanged between e16 and e16.5, however the TdTomato coverage of these MV tracks would be higher at e16.5 (**Figures 2Cii-iii**). If the latter mechanism was true, at e16.5, the length of MV tracks would be higher, and the TdTomato coverage of these MV tracks would increase or remain unchanged (**Figures 2Cv, vi**). To test these two potential mechanisms, *Cx40CreER; Rosa26^TdTomato^*; *Tie2::eGFP::Claudin5* mouse line was used, where MV tracks can be clearly visualized with eGFP reporter (Figures **2B** and **S5**). Confocal imaging of e16 (Figure **2D**) and e16.5 (Figure **2I**) whole brains, at single cell resolution allowed examination of watershed regions at both developmental time-points (Figures **2E**-**2H** and **2K-2N** respectively). Next, robust quantification of MV tracks for each watershed region was performed (Figures **2H**, **2N** and **2J**). Amongst eGFP^+^ vessels, an MV track has a distinct morphology from the surrounding capillary plexus and can be identified as eGFP^+^ tubular structures connecting the distal artery branches (Figures **2F** and **2L**). We identified an eGFP^+^ vessel as a MV track if 1) it is a continuous tubular structure and 2) is connected to a TdTomato^+^ artery at least at one end. These MV tracks were traced on multiple z-slices, using Simple Neurite Tracer (Figures **2H** and **2N**), and next, the extent of TdTomato coverage was calculated for total length of MV tracks per watershed (Figures **2H**, **2N** and **2O**). Quantifications show that while the total length of MV tracks was increased by 1.5 fold in e16.5 (Figures **2H**, **2N** and **2J**), the TdTomato coverage of these MV tracks did not change between e16 and e16.5 brains (Figures **2H**, **2N** and **2O**). These results support the second mechanism (Figures **2C**-**i→v→vi→iv**); thus, suggesting that pial collateral development is accomplished by simultaneous growth of MV tracks and artery-tips on them.

**Figure 2:**
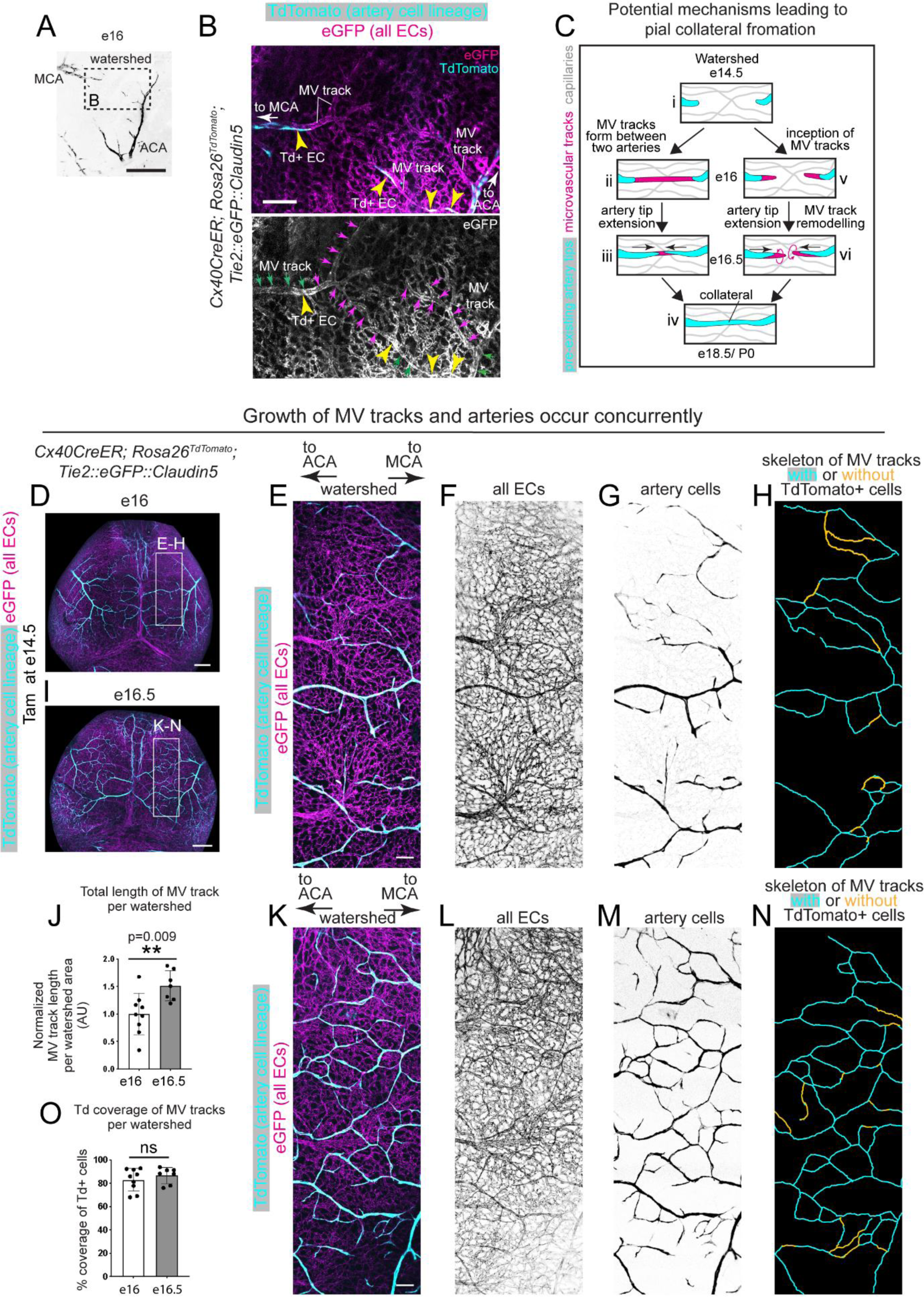
Analyses of development of MV (microvascular) tracks and artery tips during pial collateral development. (**A**) In vivo imaging of live mouse embryo at e16 shows the cerebral arteries spanning the pial watershed in mouse brain. (**B**) Inset from **A** showing TdTomato^+^ artery endothelial cells (yellow arrowheads) on eGFP^+^ microvascular tracks (arrows). Yellow arrows point to microvascular tracks partially or completely occupied by TdTomato^+^ artery ECs. Magenta arrows point to microvascular tracks at the growing front of the TdTomato^+^ labelled arteries. (**C**) Two potential mechanisms for pial collateral formation. Hypothesis 1: (i) An artery-free e14.5 watershed region (ii) is first occupied by microvascular tracks (magenta) connecting two arteries (cyan) by e16. (iii) Artery tips then start extending on pre-determined MV tracks at e16.5. Arrows indicate direction of artery tip migration. (iv) This process leads to pial collateral formation. Hypothesis 2: Alternatively, (v) while MV tracks start appearing at e16, (vi) they actively remodel and grow in length to allow artery tip extensions in a directional manner. This leads to formation of pial collaterals. Cyan represents pial artery, magenta represents MV track. (**D-N**) Concurrent MV track growth and artery tip extension. (**D and I**) Whole brain images of e16 and e16.5 embryos respectively, from *Cx40CreER; Rosa26^TdTomato^*; *Tie2::eGFP::Claudin5* mice, where TdTomato (cyan) marks pre-existing artery endothelial cells and eGFP (magenta) marks all endothelial cells. **E**-**H** is inset shown in **D**. (**E**) Higher magnification confocal image of watershed region (715 µm width) between ACA and MCA. All endothelial cells are in magenta and artery endothelial cells in cyan. (**F**) eGFP^+^ vessels in the selected watershed from e16 watershed shown in **E**. (**G**) Pre-existing artery cells in the watershed from e16 watershed shown in **E**. (**H**) Skeleton of all the eGFP^+^ microvascular tracks traced in the e16 watershed (shown in **E**) through multiple z-slices, using Simple Neurite Tracer. Traced MV tracks occupied by TdTomato^+^ artery endothelial cells are shown in cyan and MV tracks not occupied with TdTomato is shown in yellow. (**J**) Quantification of total length of microvascular tracks per watershed area (Two-tailed unpaired *t* test, p=0.009, **p < 0.01) and (**O**) Quantification of percentage TdTomato coverage of microvascular tracks in watershed regions from e16 (9 watersheds from N=9 embryos) and e16.5 (7 watersheds from N=5 embryos) brains. Data shown as mean ± standard deviation. (Two-tailed unpaired *t* test, p=0.4085, ^ns^ non-significant, p>0.05). **K**-**N** is inset shown in **I**. (**K**) Higher magnification confocal image of watershed region (715 µm width) between ACA and MCA. All endothelial cells are in magenta and artery endothelial cells in cyan. (**L**) eGFP^+^ vessels in the selected watershed from e16 watershed shown in **K**. (**M**) Pre-existing artery cells in the watershed from e16 watershed shown in **K**. (**N**) Skeleton of all the eGFP^+^ microvascular tracks traced in the e16 watershed (shown in **K**) through multiple z-slices, using Simple Neurite Tracer. Traced MV tracks occupied by TdTomato^+^ artery endothelial cells are shown in cyan and MV tracks not occupied with TdTomato is shown in yellow. Tam, Tamoxifen; MCA, Middle Cerebral Artery; ACA, Anterior Cerebral Artery; MV, Microvascular; Td, TdTomato; e, embryonic day; EC, Endothelial cell; AU, Arbitrary units. Scale bar: A, 500µm; B, 100µm; D and I, 500µm; E and K, 100µm.

### Artery ECs are the major cellular contributors of pial collaterals during embryogenesis

We next assessed the vascular-lineage composition of pial collaterals at a later stage when the collateral artery network has fully formed. This would allow us to: 1) determine if the artery cell contribution to nascent pial collaterals observed at e16.5 (Figure **1E**) persisted in later stages and 2) check contribution from microvascular lineage which is evident in newly formed collaterals in heart^10,14^ and skeletal muscle.^15^ Genetic lineage tracing with *Cx40CreER; Rosa26^TdTomato^* and *ApjCreER; Rosa26^TdTomato^* mice were performed independently, where TdTomato expression could be conditionally induced, with Tamoxifen, specifically in artery ECs or capillaries respectively. Tamoxifen was administered to timed pregnant dams at e14.5 when both MCA and ACA arteries are present (Figures **3A**, **S2G** and **S3G**). Embryos were analysed at e18.5 or P0 when the pial collateral artery network has completely formed (Figures **3A**, **S2H** and **S3H**). Whole embryonic brains were immunostained for Cx40 and segments of collateral arteries between MCA and ACA were analysed for lineage contribution (Figures **3B**-**3J**).

**Figure 3:**
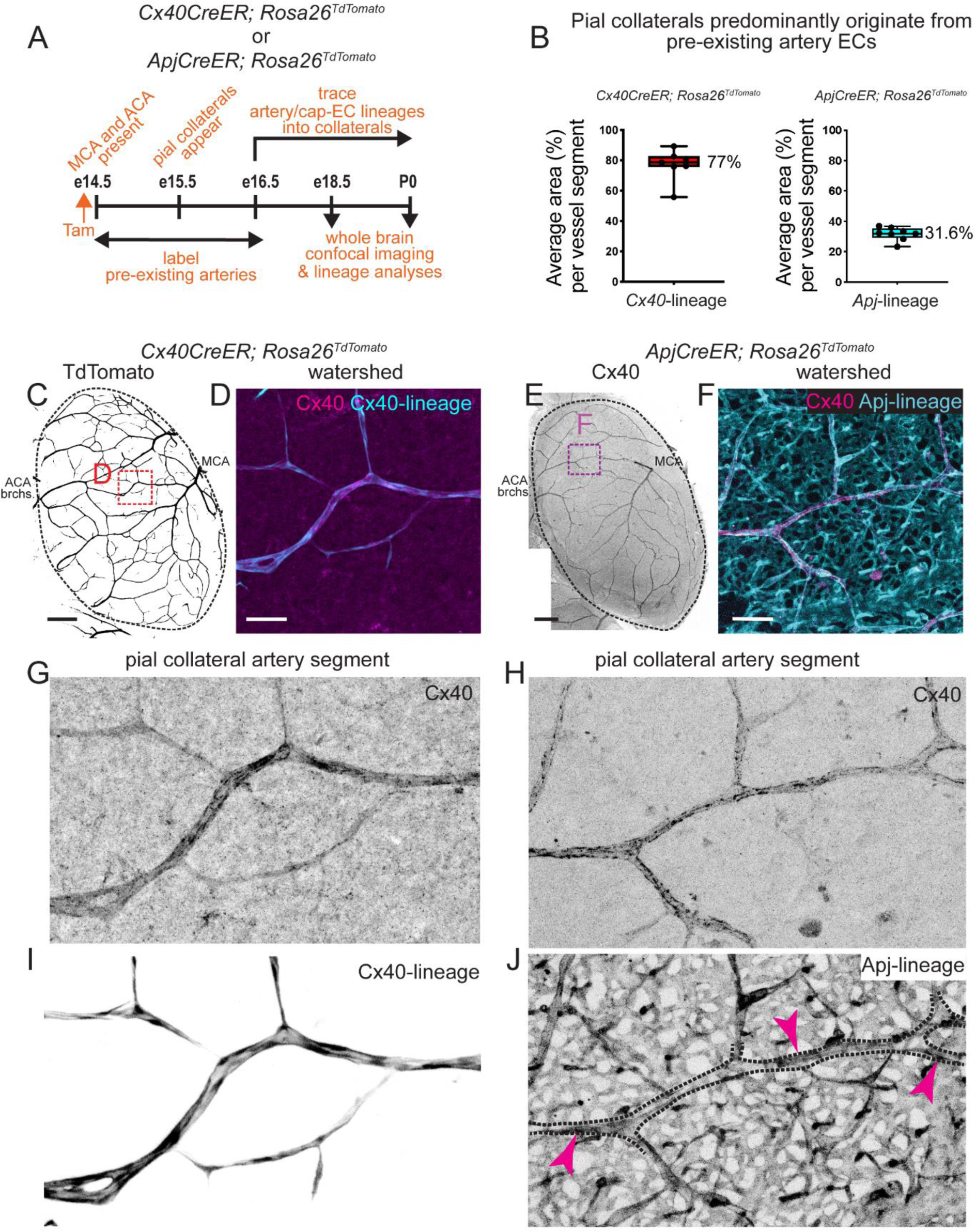
Cellular origin of pial collaterals. (**A**) Experimental design to assess lineage contribution from artery ECs and capillary ECs towards pial collaterals at e18.5/ P0. (**B**) Graph shows the average percent area of collateral segments covered with TdTomato^+^ cells in two independent genetic lineage tracing experiments using (left) *Cx40CreER* (N=7 biological replicate) and (right) *ApjCreER* (N=8 biological replicate) mouse lines respectively. The whiskers in the boxplot extends from minimum to maximum value. (**C**) Representative image of whole right hemisphere from *Cx40CreER*; *Rosa26^TdTomato^* e18.5 embryos. Black shows TdTomato^+^ arteries. (**D**) Boxed region of interest with a collateral segment from the watershed area shown in **C**. The pial collateral is lineage traced for artery ECs (cyan) and immunostained with Cx40 (magenta). (**E**) Representative image of whole right hemisphere from *ApjCreER*; *Rosa26^TdTomato^* e18.5 embryos. Black shows Cx40 immunostaining. (**F**) Boxed region of interest with a collateral segment from the watershed area shown in **E**. The pial collateral is lineage traced for capillary ECs (cyan) and immunostained with Cx40 (magenta). (**G**, **I**) Collateral segment from **D** is (**G**) immunostained for Cx40 and (**I**) lineage traced with TdTomato. (**H**, **J**) Collateral segment from **F** is (**H**) immunostained for Cx40 and (**J**) lineage traced with TdTomato. Arrowheads point to TdTomato^+^ cells from Apj^+^ lineage on the collateral segment. e, embryonic day, P, postnatal day; MCA, Middle cerebral artery; ACA, Anterior cerebral artery; brch., branch; EC, endothelial cell; Tam, Tamoxifen. Scale bars: **C** and **E**, 500µm; **D** and **F**, 100µm

We assessed the extent of lineage contribution from artery or capillary ECs towards pial collaterals in two separate experiments. As expected, all pre-existing artery ECs were labelled and traced in *Cx40CreER; Rosa26^TdTomato^* embryos (Figures **3C**, **3D**, **3G** and **3I**). Whole brain imaging of the embryo (Figure **3C**) allowed identification of pial collateral arteries present between MCA and ACA in the watershed (Figure **3D**). Individual collateral segments as identified by Cx40 immunostaining were analysed for presence of TdTomato^+^ artery ECs (Figures **3D**, **3G** and **3I**). This analysis revealed that 77% of the Cx40^+^ collateral area to be TdTomato^+^ (Figure **3B**). In contrast to this data, a similar analysis in an independent experiment with *ApjCreER; Rosa26^TdTomato^* mice (Figures **3E**, **3F**, **3H** and **3J**) showed only 31.6% of the collateral area to be TdTomato^+^, i.e. traced from capillaries (Figure **3B**). Thus, artery ECs are the major contributors of pial collateral arteries during development.

### Arterial VegfR2 mediates artery tip extension and is essential for pial collateral artery development

We analysed similarities, if any, in the molecular programs used by artery endothelial cells during collateral development in the heart and brain. Post-injury, neonatal coronary artery ECs undergo extensive migration and proliferation to build new coronary collateral arteries in the watershed. The processes of artery cell migration and proliferation are heavily dependent on Cxcr4^10^ and VegfR2^16^ respectively. To assess the functions of Cxcr4 and VegfR2 in pial collateral development, we deleted the respective genes at e14.5 and assessed the extent of pial collateral network at e18.5/P0 by whole brain imaging (Figure **4A**). Surprisingly, deletion of arterial *Cxcr4* did not affect the pial collateral network at e18.5 (Figures **4B****, S7A** and **S7B**) or in adulthood (Figures **S7C**-**S7E**). This data suggests that a Cxcr4-independent vascular mechanism facilitates pial collateral development and its maintenance.

**Figure 4:**
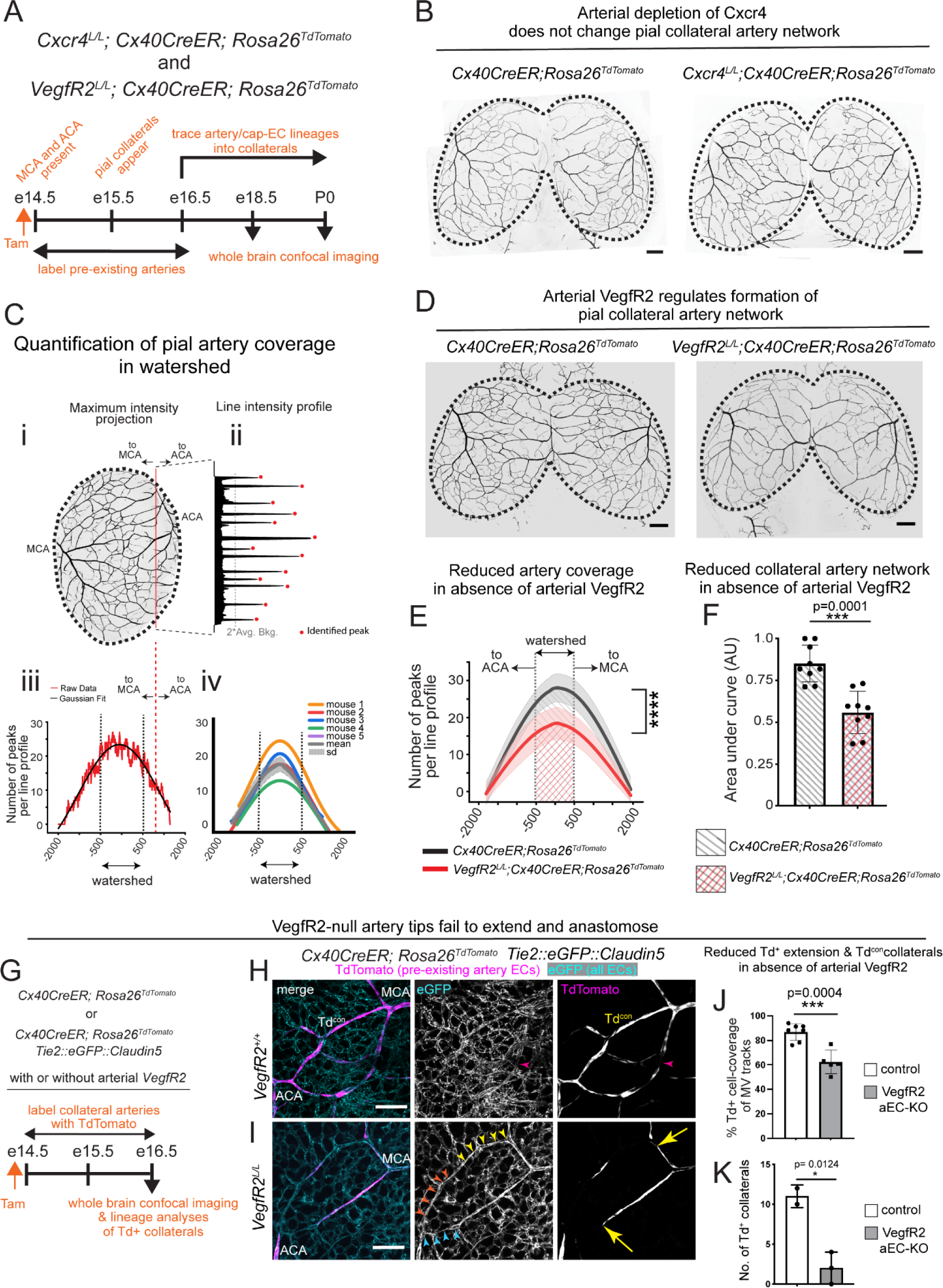
VegfR2-dependent formation of pial collateral arteries. (**A**) Experimental design to lineage label and trace pre-existing artery endothelial cells in pial layer, with or without arterial *VegfR2* and *Cxcr4* at e18.5/ P0. (**B**) Representative confocal images of whole brains with and without arterial *Cxcr4*. Black shows pre-existing artery endothelial cells. Control, N= 8 and artery specific knockout for *Cxcr4*, N= 13. (**C**) Summary of method to quantify pial artery coverage across the entire brain including watershed. (**D**) Confocal images of whole brains with and without arterial *VegfR2*. Black shows pre-existing artery endothelial cells. (**E**) Quantification of overall pial artery coverage with and without arterial *VegfR2* quantified using method shown in **C**. KS test performed on the distributions generated D value= 0.569, p value= 2.2066 X 10^-141^. (**F**) Quantification of pial collateral artery network with and without arterial *VegfR2* quantified using method shown in **C**. N=9 biological replicates for each group. Two-tailed unpaired *t* test, p=0.0001. (**G**) Experimental design to assess the effect of *VegfR2* deletion from arteries at e16.5. (**H**, **I**) Confocal images of pial watershed region showing TdTomato-contiguous pial collateral arteries (Td^con^) in magenta and microvasculature in cyan. Arrowheads show part of microvascular track not occupied by TdTomato^+^ artery cell, arrows point to artery tips on microvascular tracks. (**J**) Quantification of TdTomato^+^ artery cell coverage of pre-existing MV tracks with (N=7 biological replicates) or without (N=5 biological replicates) arterial *VegfR2*. Two-tailed unpaired *t* test, p=0.0004. (**K**) Quantification of pial collateral arteries contiguously populated with TdTomato^+^ artery cells, with (N=2) and without (N=3 individual brains) arterial *VegfR2*. Two-tailed unpaired *t* test, p=0.0124. Each dot in F, J, and K represents a biological replicate. Data shown as mean ± standard deviation. *p < 0.05 and ***p < 0.001. MCA, Middle cerebral artery; ACA, Anterior cerebral artery; e, embryonic day; P, postnatal day; Tam, Tamoxifen; EC, endothelial cell; Td^con^, TdTomato-contiguous; AU, Arbitrary Unit; aEC, artery endothelial cell; KO, knockout. Scale bars: **B**, **D**, 500µm; **H**, **I**, 100µm.

We next assessed if pial collateral artery development is dependent on arterial VegfR2. Whole brain imaging of control and *VegfR2* artery specific knockout embryos showed differences in collateral connectivity in the watershed, and required systematic quantification (Figures **4C**-**4F**). The extensive connectivity of pial arteries across the hemisphere (including watershed) provides several possible collateral paths for the blood to flow. These artery-artery connections are innumerable. We developed a method that allowed us to compare the pial artery coverage between the control and *VegfR2* artery EC specific knockout brains quantitatively (Figure **4C**). Intensity line-profiles were generated across the width of the hemisphere (Figures **4C i, ii**). The identified peaks per line-profile was plotted across the width of the hemisphere (Figure **4C iii**). The data was fit to a Gaussian function and the fitted curves were used as representative of the arterial coverage per hemisphere in each mouse (Figures **4C iv**). The average of the fitted curves represented one genotype (grey curve, Figure **4C iv**). Quantification of pial artery coverage showed significantly different distributions between control and *VegfR2* knockout brains (KS test, D value=0.569, p value=2.2066 X 10^-141^) (Figure **4E**). Peak arterial coverage was observed at the centre of the hemisphere and watershed was defined as a region ± 500µm from the centre (Figure **4E**). The loss of arterial coverage in the watershed was evident in the knockout (Figure **4E**) and was quantified as area under the curve (Figures **4E** and **4F**). This area under the curve represented the extent of collateral artery network in the watershed and was significantly lower in brains without arterial *VegfR2* (Figure **4F**). Thus, depletion of *VegfR2* from arteries reduced the extent of pial collaterals formed during embryogenesis.

We next investigated the possible mechanisms by which arterial VegfR2 facilitated pial collateral formation. *VegfR2* was deleted from artery ECs at e14.5, and analyses of pial collateral network was performed at e16.5 (Figure **4G**), a timepoint when collaterals are beginning to form via artery tip extensions (Figures **1D**-**1G**). A thorough analyses of the pial watershed at e16.5 was performed to assess the extent of MV tracks and TdTomato (artery-lineage) coverage (Figures **S8A** and **S8B**). Close analyses of collaterals in control brains showed that TdTomato^+^ artery cells occupied a major length of microvascular tracks connecting branches of MCA and ACA (Figure **4H**). In contrast to the control brains, VegfR2-depleted brains showed reduced TdTomato^+^ artery cell coverage of the microvascular tracks (Figures **4I****, 4J,** and **S8B**). Of note, without arterial VegfR2, the length of MV tracks also reduced (Figure **S8C**), without affecting the density of capillaries (Figures **S8D**, **S8E**). Quantification of collateral arteries showed fewer TdTomato contiguous or Td^con^ collateral arteries, without arterial VegfR2 (Figure **4K**). This indicated that initiation of collateral development at an earlier timepoint, e16.5, also depends on arterial VegfR2. This VegfR2-dependent collateral phenotype (Figures **4J** and **4K**) was independent of artery cell proliferation which was minimal at this stage, for both control and VegfR2-deleted brains (data not shown). Interestingly, this was in contrast to the cell cycle state of ischemic coronaries, which is 10-fold higher than pial arteries.^10,16^ Together, arterial VegfR2 regulates pial collateral formation by promoting artery extensions along microvascular tracks.

We next investigated if the presence of microvascular cells on pial collaterals (Figure **3B**) is a result of arterialization, i.e., differentiation of capillary cells into artery endothelial cells. To address this question, we assessed the potential role of Dach1, a transcriptional factor, which drives arterialization of cardiac capillary cells into coronary artery endothelial cells, during development and post-injury.^17^ As observed in developing mouse hearts,^18,19^ expression of Dach1 was higher in pial capillaries as compared to pial arteries (Figure **S9A**). Next, *Dach1* was deleted from capillaries using *ApjCreER* at e14.5, when both cerebral arteries are present but a pial collateral network is absent. Whole brains were then analysed for pial collateral network at e18.5 (Figure **S9B**). We observed a robust deletion of *Dach1* from capillary ECs (Figures **S9C**-**S9D**). However, no significant difference was observed in the number of Cx40^+^ collaterals in the *Dach1* capillary-EC specific knockout brains (Figure **S9E**). Thus, in contrast to coronary artery development,^19^ the formation of pial collaterals during embryogenesis is independent of capillary Dach1 activity.

### Pial collaterals do not remodel in adulthood

We asked the question if arterial VegfR2 could modulate neonatal pruning events. To address this, *VegfR2* was deleted in arteries at P0, and whole brain imaging was performed at P2, P7 and P14 (Figure **S10A**). The arterial deletion of VegfR2 was robust at P7 (Figures **S10B**-**S10C**). Analyses of whole P2 (Figures **S10D**-**S10G**), P7 (Figures **S10H**-**S10K**) and P14 (Figures **S10L**-**S10O**) hemispheres revealed no significant differences in arterial coverage with depletion of arterial VegfR2 (Figures **S10G, S10K** and **S10O**). Thus, arterial VegfR2 is dispensable for neonatal pial artery and collateral artery pruning.

We next investigated the cellular dynamics associated with fully formed and mature pial collateral arteries between MCA and ACA in adult mice. For this, we utilized an extensive longitudinal in vivo imaging paradigm that allowed us to get single cell resolution of vascular structures. 3-5 months old *Cx40CreER; Rosa26^TdTomato^*; *Tie2::eGFP::Claudin5* mice underwent craniotomy and an optical window of 4mm was implanted on one of the hemispheres between the cranial landmarks—bregma and lambda (Figures **5A** and **5B**). In these transgenic mice, pre-existing pial arteries were labelled with TdTomato upon Tamoxifen administration (Figure **5C**), and constitutively expressing eGFP marked all endothelial cells. 5 consecutive doses of Tamoxifen resulted in 95.4% and 86.8% recombination in proximal and distal arteries respectively, but a lower 65.3% recombination in the pial collateral regions (Figures **5D** and **S11**). A lower recombination efficiency in the pial collateral arteries resulted in a patchy reporter pattern comprising of a mixture of TdTomato^+^ and TdTomato^-^ cells (Figure **5D**). Like in developmental stages (Figure **3B**), artery ECs populate adult pial collaterals to a greater extent as compared to other vascular lineages. This ultimately allowed us to most accurately track the position of individual cells within a given collateral artery.

**Figure 5:**
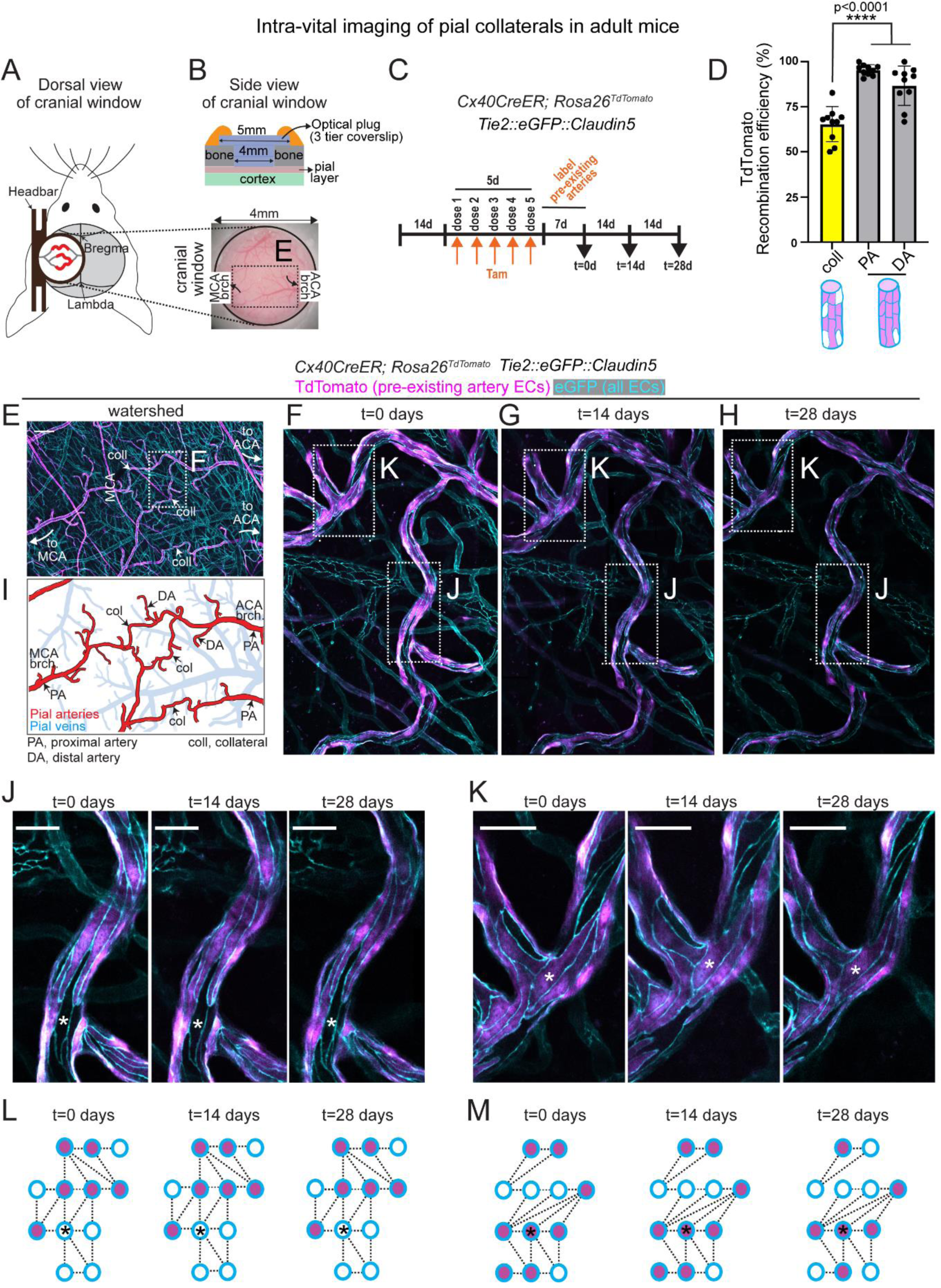
Longitudinal live-imaging of adult pial collaterals. (**A**, **left**) Dorsal view of the cranial window created with the attached headbar on adult mouse head. (**A**, **right**) Brightfield image of the 4mm cranial window obtained with a stereoscope shows the pial vasculature. Dotted rectangle encompasses MCA, ACA and the watershed between them and is imaged at high resolution as shown in **E**. (**B**) Schematic of the side view of the cranial window showing the 3-tier optical plug inserted into the skull giving access to pial layer. (**C**) Experimental design for longitudinal imaging of pial vasculature in adult mice. (**D**) Quantification of TdTomato reporter recombination in pial collateral arteries and different segments of cerebral arteries. Each dot represents individual vessel. One-way ANOVA was done to test significance statistically (Tukey’s mean comparison test, PA vs. collateral, p<0.0001; DA vs. collateral, p<0.0001; PA vs. DA, p=0.08). ****p < 0.0001. (**E**-**K**) Images of pial watershed with pre-existing artery endothelial cells labelled in TdTomato (magenta) and all endothelial cells labelled in eGFP (cyan). (**E**) Confocal image from optical window created for longitudinal imaging of *Cx40CreER; Rosa26^TdTomato^; Tie2::eGFP::Claudin5* mouse as shown in Figure 4A. The image shows both pial and dural vessels. (**F**-**K**) Confocal images of pial collateral arteries from cranial window shown in **E**, across 3 timepoints, each 2 weeks apart. (**I**) Pial layer segmented out from **E**. red and blue show pial arteries and veins respectively. **J**, **K** are the insets shown in **F**-**H**. (**L**, **M**) Cellular distribution of TdTomato^+^ artery cells and eGFP^+^ endothelial cells in collateral artery segments shown in **J** and **K** respectively. * represents a reference cell. MCA, middle cerebral artery; ACA, anterior cerebral artery; d, days; Tam, Tamoxifen; PA, proximal artery; DA, distal artery; coll, collateral; brch., branch; EC, endothelial cell. Scale bars: **E**, 200µm; **J**, **K** 25µm

An optical window allows longitudinal imaging. Imaging the collateral zone (watershed between MCA and ACA) for a month (Figure **5E**), resulted in high quality images with single cell resolution (Figures **5E**-**5H**). Morphology of the pial vessels alongside reporter activity in endothelial cells were instrumental in identifying pial arteries, veins and capillaries (Figures **5E** and **5I**). High resolution images of artery endothelial cells (both TdTomato^+^ eGFP^+^ and TdTomato^-^ eGFP^+^) within a collateral artery segment across many collaterals and optical windows were analysed (Figures **5J** and **5K**). Data showed that artery endothelial cells do not remodel or reposition themselves within a given collateral artery segment (Figures **5J**-**5M**). Cellular matrices representative of the relative position of collateral artery cells suggested that under physiological conditions, every artery cell within a mature/ adult pial collateral maintains its contact with the same neighbouring cells (Figures **5L** and **5M**). Thus, surprisingly, the cellular organization within pial collaterals and the overall collateral structure was extremely stable. Together, our data with cellular resolution, demonstrate the static nature of artery endothelial cells during homeostasis.

To investigate if cells of pial collaterals reposition in response to hypoxic conditions, we next subjected the mice to 7% atmospheric oxygen. We utilized a similar imaging paradigm as shown in Figure **5**. Longitudinal imaging of cranial windows allowed visualization of pial vasculature at cellular resolution (Figures **S12A**-**S12B**). *Cx40CreER; Rosa26^TdTomato^; Tie2::eGFP::Claudin5* mice were used to label/trace pre-existing arteries, followed by exposure to low (7%) atmospheric oxygen (Figure **S12C**). High resolution long-term imaging of every cranial window revealed that diameter of collateral arteries between MCA and ACA increased with low tissue oxygen (Figures **S12D**-**S12F**). This increase in diameter was not due to an increase in artery cell number (Figure **S12G**), but because of an increase in size of individual artery cell (Figures **S12H**-**S12J**). Together, artery endothelial cells within pial collateral arteries do not reorganize, even under hypoxic conditions.

### Arterial VegfR2 is dispensable for pial collateral maintenance

We next asked if arterial VegfR2 is necessary for the maintenance of adult pial collaterals using adult mice deficient in arterial *VegfR2*. Using the same in vivo imaging paradigm mentioned above, pial collateral segments in artery endothelial cell specific *VegfR2* knockout mice were imaged at cellular resolution and followed for a month with imaging time-points separated by 2 weeks (Figure **6A**). Longitudinal imaging through an optical window (Figure **6B**) yielded a map of vascular network which could be followed for months (Figures **6C** and **6D**). Of note, our qualitative observation suggests that *VegfR2* depleted pial collaterals had smaller diameters. High magnification images of watershed between MCA and ACA (Figures **6E**-**6G**) allowed us to analyse individual *VegfR2*^-/-^ artery endothelial cells (TdTomato^+^) residing on adult collateral arteries (Figures **6H**-**6J**). Similar to wild-type (Figure **S11**), *VegfR2* deleted proximal arteries showed very high TdTomato recombination (Figure **S13A**). Long term imaging of the same collateral regions across many collaterals and mice showed no significant changes in cellular positions relative to their neighbouring cells (Figures **6K**-**6M** and **S13B**). Thus, arterial VegfR2 is dispensable for maintenance of mature pial collateral arteries. Together, in this study, we highlight the essential role of arterial VegfR2 in promoting artery tip extension and pial collateral development during embryogenesis (Figure **6N**); a function dispensable in adulthood.

**Figure 6:**
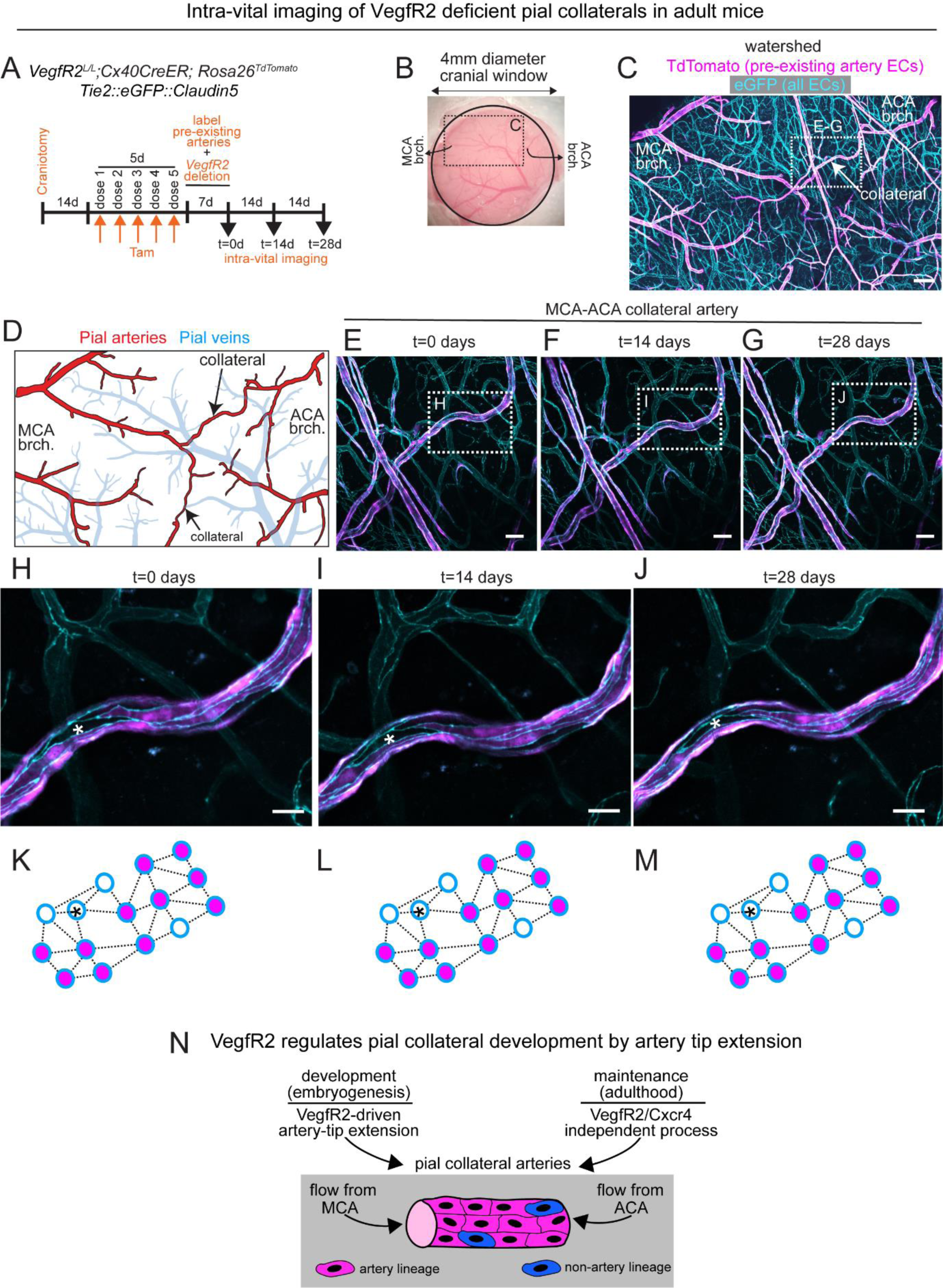
Analyses of role of arterial VegfR2 in pial collateral maintenance. **(A)** Experimental design to visualize artery cell dynamics in pial layer of adult mice without arterial *VegfR2*. (**B**) Brightfield image of cranial window showing pial vasculature. The dotted rectangle encompasses branches of MCA, ACA and the watershed between them. (**C**) Confocal image of cranial window created in mice without arterial *VegfR2* as shown in **B**. Magenta shows all artery cells, cyan shows all endothelial cells. (**D**) Pial layer segmented out from **C** shows pial arteries and veins in red and blue respectively. (**E**-**G**) Confocal images of collateral artery from three timepoints, each 2 weeks apart as shown in the boxed region in **C**. (**H**-**J**) Confocal images of collateral artery segments shown in **E**-**G**. (**K**-**M**) Cellular distribution of TdTomato^+^ artery cells and eGFP^+^ endothelial cells in collateral artery segments shown in **H**-**J**. * represents a reference cell. (**N**) Schematic summarizing the role of arterial VegfR2 in development and maintenance of pial collateral arteries. While arterial VegfR2 facilitates tip extension of pre-existing arteries during development, it is dispensable for pial collateral maintenance in adulthood. Pial collaterals, both in developing and mature brains, are majorly populated by endothelial cells of artery-lineage. MCA, middle cerebral artery; ACA, anterior cerebral artery; d, days; Tam, Tamoxifen; EC, endothelial cell; brch., branch. Scale bars: **C**, 150 µm; **E**-**G**, 50µm; **H**-**J**, 20µm.

## DISCUSSION

In the present study, we investigate artery cell dynamics during collateral artery formation and maintenance in mouse brain. Firstly, using an artery-specific molecular marker, we dissect out where and when pial collaterals appear during embryogenesis. Secondly, we develop a live imaging method in mouse embryos to provide conclusive evidence that pre-existing arteries grow on pre-determined microvascular tracks which then connect to build pial collaterals *de novo*. Thirdly, by combining mouse genetics with whole brain confocal imaging at cellular resolution, we show that an angiogenic vascular receptor, VegfR2, facilitates artery-tip extension on microvascular tracks and aids origin of pial collaterals. Finally, we show that Cxcr4, a receptor essential for artery reassembly and formation of coronary collateral arteries *de novo*, is dispensable for development of pial collaterals. Thus, two fundamentally distinct mechanisms contribute to the origin of coronary and pial collaterals.

### Organ-specific mechanisms build collaterals

In this study, we provide direct support that a vascular mechanism driven specifically by artery endothelial cells leads to the origin of pial collaterals. Like in the heart, this artery specific process is regulated by arterial VegfR2, but is distinct from artery reassembly, where pre-existing artery cells dissociate from resident coronaries, migrate, de-differentiate/proliferate and coalesce to build coronary collaterals.^10,16^ Instead, developing pial arteries extend along microvascular tracks and connect with other artery tips to build new collateral artery paths. Furthermore, in heart, coronary collateral formation is regulated by arterial Cxcr4 which drives artery cell migration during artery reassembly. In contrast, arterial Cxcr4 had no impact on pial collateral formation or its maintenance. Unlike heart, pial artery endothelial cells contribute minimally via proliferation. Evidence for negligible artery cell proliferation within the pre-existing pial arteries prompts us to hypothesize that formation of pial collaterals during embryogenesis is minimally dependent on proliferation of pre-existing artery ECs. That being said, the underlying mechanism leading to extension of distal artery tips remain unclear. It is possible that distal artery endothelial cells experience a forward push from the proximal segments of the artery. This forward pressure could be generated due to incorporation of new cells. Accommodating these additional cells within an artery would require further cellular organization throughout the length of the artery, and could lead to distal extension of the artery tips. Finally, the impact of blood flow on cellular organization within pre-existing arteries and/or its extension, remains an open question for future investigation.

The need for functional collaterals is most during an ischemic condition such as an artery occlusion, and can significantly impact tissue function. The differences in factors which regulate origin of coronary and pial collaterals, are potentially driven by the metabolic needs of these tissues. While cardiomyocytes can survive hypoxia for few hours,^20^ neurons are more susceptible to low levels of oxygen.^21^ Higher hypoxia sensitivity of brain tissue may lead to formation of pial collaterals during embryogenesis as a protective measure, while coronary collaterals form only upon induction of an arterial occlusion. Furthermore, collateral arteries in skeletal muscle,^15,22^ intestine,^23^ and hind limb^24^ can also undergo arteriogenesis.^25^ Identifying collateral-building vascular mechanisms across organs could reveal clinically relevant information and open avenues for therapeutics.

### Arteries and capillaries collaborate to build collaterals

We demonstrate the formation of collateral arteries, in real time, from pre-existing arterial lineage as observed^10,11,16,26^ or proposed^7^ earlier. While this manuscript was in progress, Perovic et al reported that two separate vascular lineages populate pial collaterals.^11^ We also observed that well-formed pial collaterals are majorly populated by artery cells, and minimally by capillaries. Intra-vital imaging of embryonic brains reveal that pre-existing artery cells extend along defined microvascular paths to connect and build pial collaterals. These artery specific events are executed within a short span, are difficult to capture using static images and can be observed only by live imaging.

We cannot exclude the contribution of other artery-driven mechanisms. We observed single recombined artery cells migrating towards pre-existing arteries; coalescence of these cells with arteries could lead to arterial growth. Alternatively, the migration and incorporation of single artery cells at the proximal ends of the artery could indirectly contribute towards distal elongation of arteries. The dynamics of single artery cell movements could also establish the branching pattern of the pial artery network; the precise mechanisms for which are unknown.

We show that pre-existing artery tip extensions occur along a distinct supporting structure. In the heart, single artery endothelial cells migrate by hitch-hiking on to the capillary networks during the formation of new collaterals. On the contrary, our study in the embryonic brains reveals the presence of a unique (Tie2^+^) vascular segment on which the arteries extend. These microvascular tracks are distinct from the surrounding capillary mesh in that they anastomose ACA and MCA arterial branches but do not show any artery specification. We speculate that the microvascular tracks form during the specific time window (e16-e17) which coincides with the period when majority of collaterals appear. Thus, these tracks may play a crucial role in pial collateral formation. Based on our Apj lineage data, we hypothesize that the cellular origin of these microvascular tracks is of capillary in nature. However, we cannot completely exclude the possibility that some tracks (guiding vessels for artery tip extensions) could also originate from mixed lineages. Thus, though capillaries do not massively contribute to the ECs of the pial collateral, they could create/remodel into MV tracks, to indirectly facilitate development of pial collaterals. Elucidating the underlying mechanism governing MV track formation is an exciting avenue for future investigation. Additionally, determining their precise molecular identity will allow creating genetic tools to assess if MV tracks are indispensable for development of pial collaterals.

Our data about pre-defined microvascular tracks guiding the growth of pial arteries suggest that the microvasculature could shape the artery network of the brain. However, the microvascular contribution towards pial collaterals via arterialization is less likely, as deletion of *Dach1*, a regulator of arterialization in heart,^17,19^ had minimal effect on pial collateral network during embryogenesis. This data contrasts the results from heart studies where deletion of *Dach1* from all endothelial cells using the *Tie2Cre* driver, had a massive impact on coronary artery development during mouse embryogenesis.^19^ Thus, we propose that as artery tips extend and get connected in the watershed, microvascular cells are “trapped” within pial collaterals. This could generate a mixed lineage distribution^11^ within pial collaterals—both in embryos and adults. Together, multiple mechanisms could be actively building pial vasculature during embryogenesis. That being said, possible contribution from endothelial progenitor cells or peri-vascular cell types (pericytes, smooth muscle cells) towards pial collateral formation cannot be entirely excluded.

### Distinct mechanism for collateral maintenance

We observed Cx40^-^ ECs co-existed with Cx40^+^ artery cells within a single adult collateral segment. The source of Cx40^-^ cells within a mature pial collateral is unclear. Our data explicitly show that unlike pial capillaries, artery ECs (Cx40^+^ or Cx40^-^) within adult pial collaterals preserve their cellular location and appear to be static structures, at least during homeostasis. Thus, it is less likely that (Cx40^-^) microvascular cells such as capillary ECs would get incorporated into the collaterals. We speculate that the (TdTomato^-^) Cx40^-^ cells are artery ECs of a different molecular origin and/or function. Determining the source or need for such molecular heterogeneity could be key to identifying special arterial roles in context of injury such as hypoxia or cerebral artery occlusions.

Pial collateral arteries undergo outward remodelling when subjected to hypoxia or ischemia.^27^ Studies show that artery endothelial cells of pial collaterals proliferate to support this outward remodelling.^11,27^ In contrast, our longitudinal imaging data provide evidence that the number (or position) of artery cells within a pial collateral segment does not change in response to hypoxia. It is possible that a longer exposure to hypoxia or if allowed to recover from these hypoxic conditions, the pial collateral artery cells would undergo proliferation leading to an increase in collateral diameter. Thus, cellular response to ischemia or hypoxia could be a multi-layered phenomenon, involving more than one mechanism.

Given that collateral arteries have a unique hemodynamic niche,^28^ it would be interesting to investigate which molecular programs maintain or remodel pial collaterals in an adult brain. During embryogenesis, pial collaterals are built *de novo*, in an artery-free zone and remodel as and when blood flows through them. In contrast, mature collaterals remodel in response to differential blood flow patterns and oxygen demands of their surrounding tissue. Postnatal remodelling of pial collaterals is dependent on recruitment of smooth muscle cells and exhibits tortuous morphology.^28^ These fundamental differences could explain the requirement for distinct molecular programs for building new collaterals and maintaining old ones. This is evident in our study; VegfR2 controlled pial artery network during embryogenesis but was dispensable in adulthood. That being said, arterial VegfR2 may regulate other aspects of vascular changes those require activation of angiogenic pathways,^29^ such as remodelling induced by hypoxia, ischemia or aging. We did not notice any remodelling events within adult pial collaterals, in presence or absence of arterial Cxcr4. Together, adult pial collaterals are maintained via mechanisms independent of two key angiogenic receptors in arteries—VegfR2 and Cxcr4. It will be interesting to test if adult collateral maintenance is independent of signals originating from an arterial niche. It is possible that surrounding microvasculature plays a more prominent role in maintenance of pial arteries and collaterals in adults. Investigating the molecular identity of the microvasculature surrounding pial collaterals could reveal new vascular mechanisms key to collateral maintenance.

## LIMITATIONS OF THE STUDY

The experimental designs of this study do not allow us to exclude the contribution of other cellular mechanisms associated with pial collateral formation during embryogenesis. Additionally, we are unable to comment on the molecular nature of artery cell heterogeneity observed in adult pial collaterals and its role (if any) on recovery from ischemia.

## MATERIALS AND METHOD

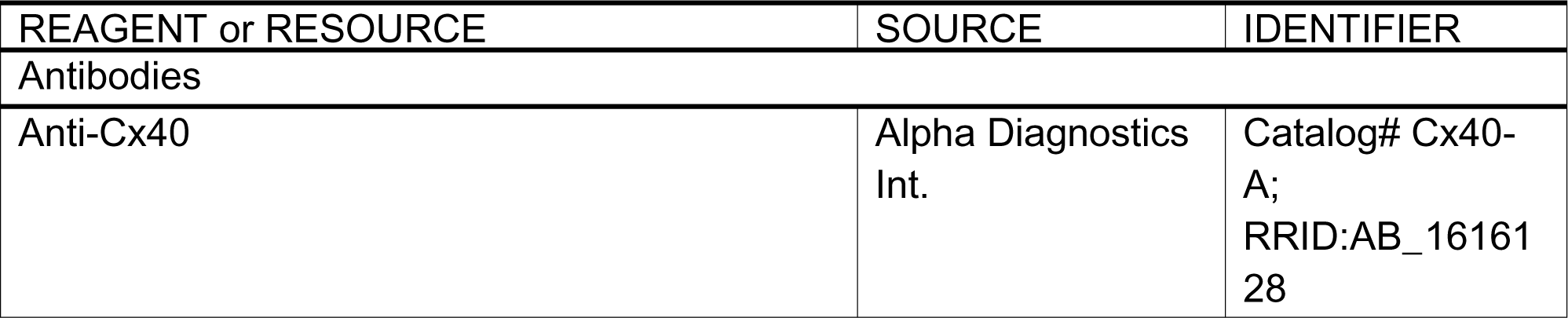
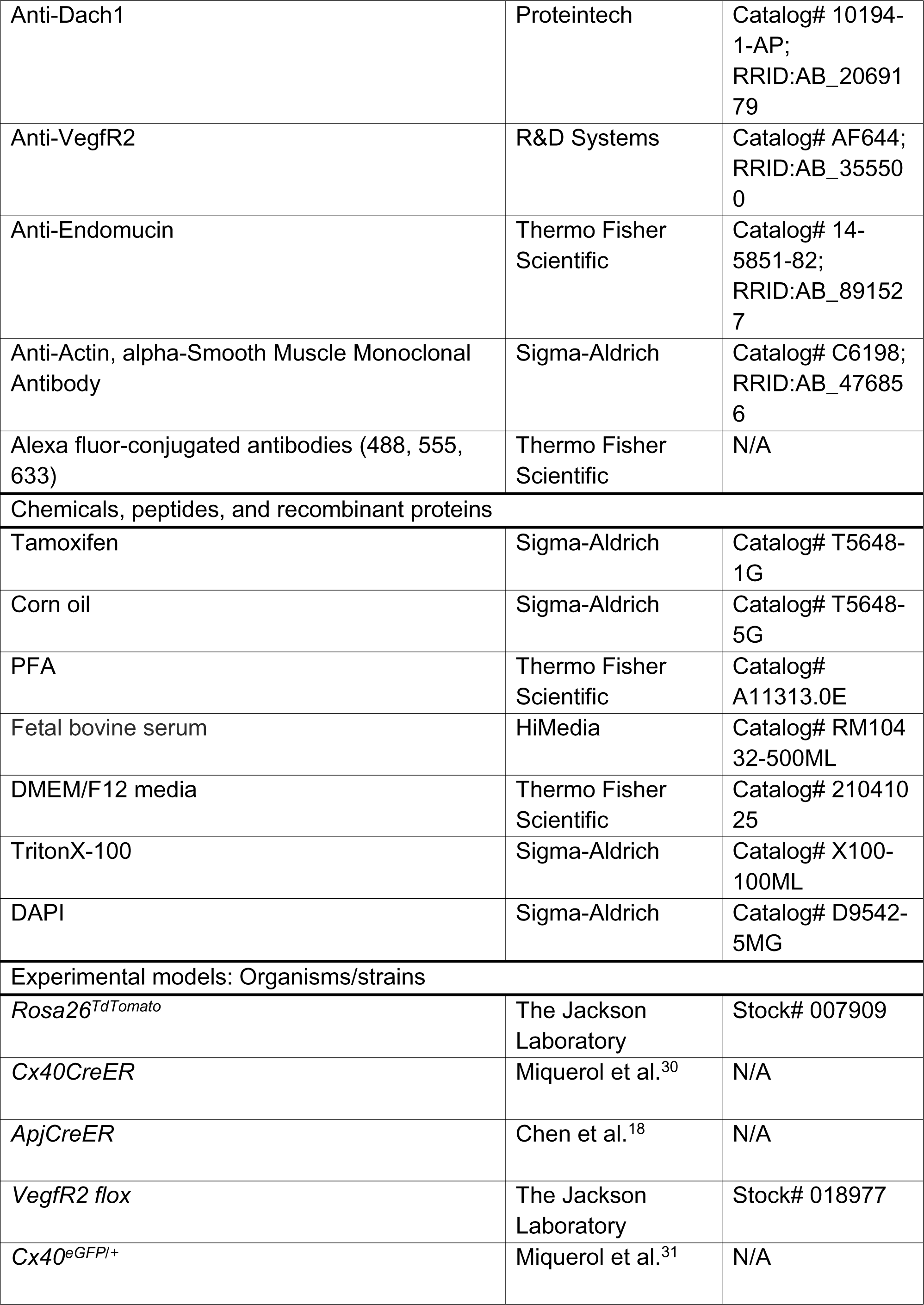
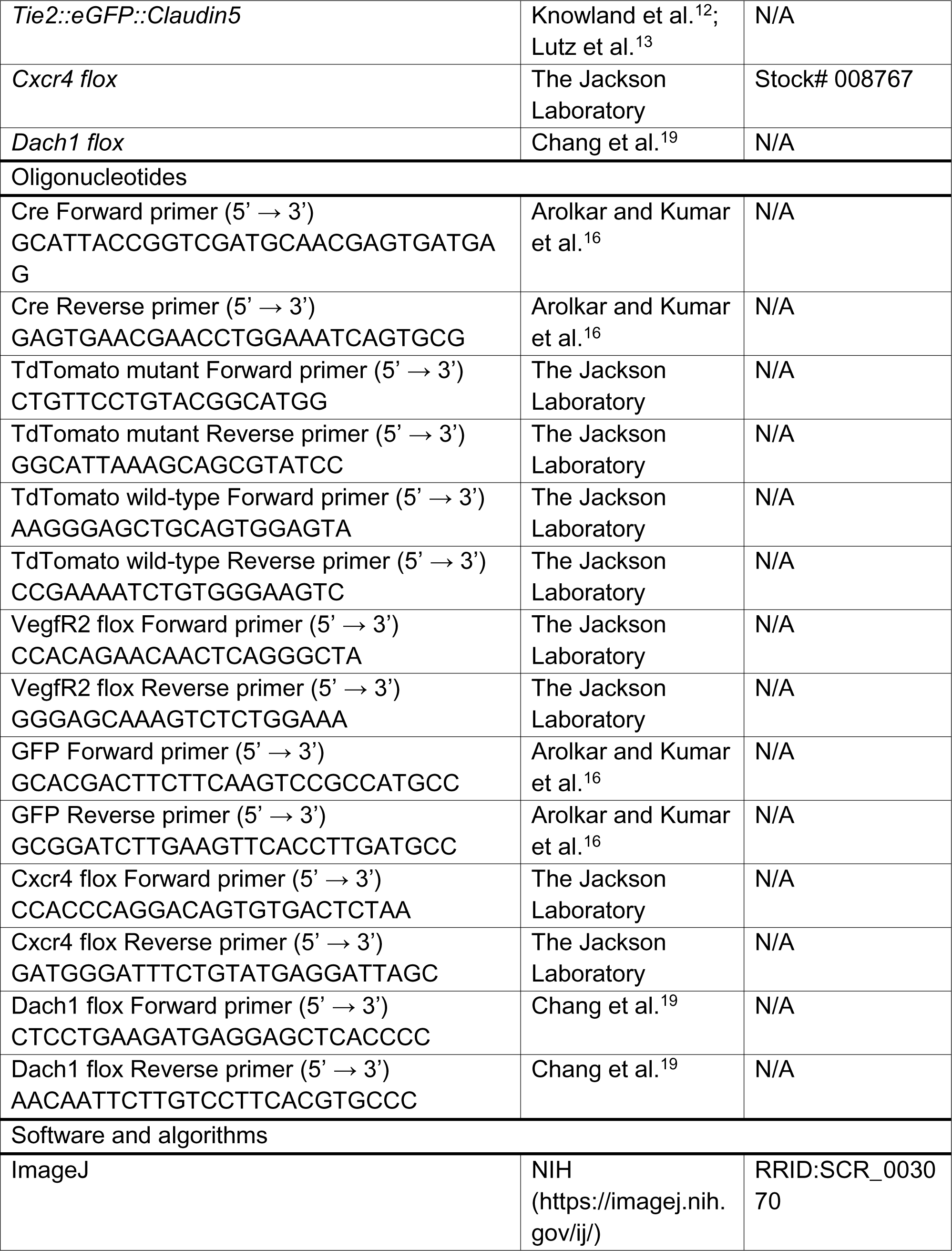
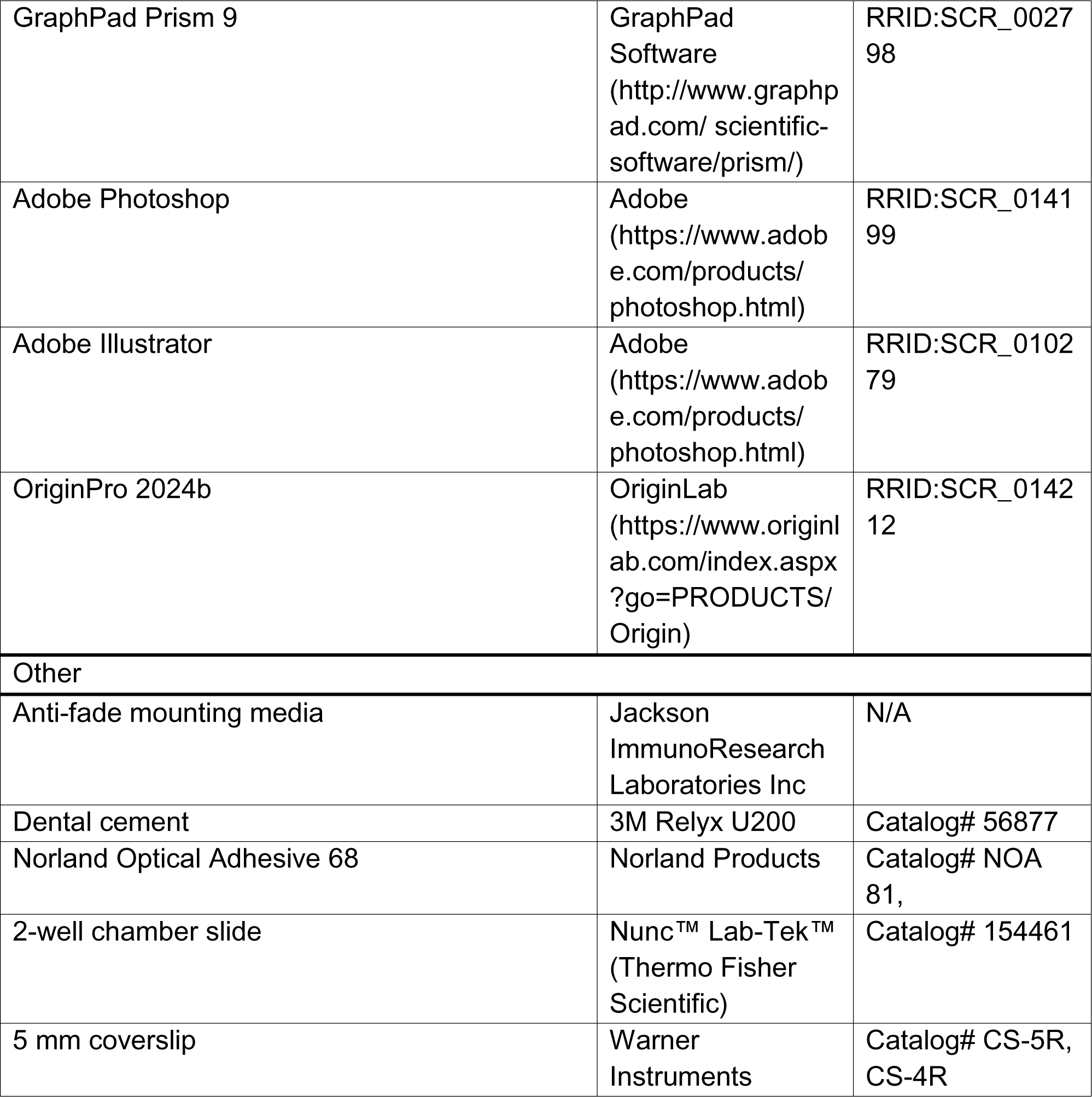

## RESOURCE AVAILABILITY

### Lead contact

Further information and requests for resources and reagents should be directed to and will be fulfilled by the lead contact, Soumyashree Das.

### Materials availability

This study did not generate new unique reagents.

### Data and code availability

- All data reported in the paper will be made available by the lead contact upon reasonable request.
- This paper does not report original code.
- Any additional information required to reanalyse the data reported in this paper is available from the lead contact upon request.

## EXPERIMENTAL MODEL AND STUDY PARTICIPANT DETAILS

### Mice

All experiments were performed on mice with mixed strain background. All mice were housed and bred at the National Centre for Biological Sciences animal facility in accordance with the Institutional Animal Ethics Committee (IAEC). All procedures were performed with approval from IAEC. Data shown in this study is compiled from age-matched males and females from several litters.

### METHOD DETAILS

### Visualization of pial arteries in fixed samples

#### 1. Harvesting whole brains/heads from mouse embryos

Depending on the developmental stages and size of the embryos, pial collateral arteries were visualized either in whole brains (e18.5 or P0) or using whole heads (e12.5-e16.5).

e12.5-e15.5: To perform the whole head explant at e12.5-e15.5 stages, the pregnant females were euthanized through carbon dioxide overdose and a Y-shape incision was made on the abdomen. Intact embryonic sacs were dissected out. Small punctures were made on the embryo sac with the help of microforceps. The whole embryo sac was fixed in 4% Paraformaldehyde (PFA) for 2 hours with gentle shaking at 4°C. Following fixation, the embryo sac was washed at 4°C with gentle shaking. This washing step was performed with 1X PBS and repeated 4 times every 15-20 mins. The embryos were then extracted from the embryo sacs and the heads were removed with the help of microforceps.

e16-e16.5: The embryonic explants at e16/e16.5 are performed similarly. However, for *Tie2::eGFP::Claudin5* genotype, the loose skin on top of the brain was removed to aid immunostaining. Whole heads were imaged within 4 hrs, prior to PFA fixation to avoid loss of GFP fluorescence signal.

e18.5: To perform the whole brain explant at e18.5 stage, the timed pregnant females were euthanized as above. The skull of each embryo was gently removed using the microforceps. The embryonic brains were scooped out with the help of a microspatula and fixed in 4% PFA at 4°C for 2.5 hours. Agitation was avoided at this stage to preserve tissue integrity of embryonic brains. The fixed embryonic brains were then washed with 1X PBS three times at 15 minutes intervals with gentle shaking at 4°C.

P0: Neonatal explants at P0 were performed by euthanizing the pups through carbon dioxide overdose followed by decapitation. The skull was removed using scissors and microforceps. The brains were scooped out using the microspatula. The brains were then fixed with 4% PFA at 4°C for 2 hrs followed by three washes with 1X PBS at an interval of 15 mins.

#### 2. Whole brain/head immunostaining of mouse embryos

e12.5-e15.5: The embryonic heads (e12.5 - e15.5) were incubated in 0.1% PBST (0.1% TritonX-100 in 1X PBS) for 2.5 hours to permeabilize the tissue, followed by incubation with primary antibodies overnight at 4°C with gentle shaking. Heads were then washed with 1X PBS, changing the buffer every 1-2 hours for 1 day. This step was followed by incubation with secondary antibodies made in 0.1% PBST. Samples were then washed every 1-2 hours for 1 day with 1X PBS. Following immunostaining the brains are immersed in an anti-fade mounting media (from Jackson ImmunoResearch Laboratories Inc.) and imaged immediately.

The embryonic and neonatal brains were incubated with primary antibody (made in 1X PBS) overnight, at 4°C with gentle shaking. Samples were then washed with 1X PBS, changing the buffer every 1-2 hours for 1 day and incubated with Alexa Fluor conjugated secondary antibodies (made in 1X PBS). Finally, samples were washed every 1-2 hours for 1 day with 1X PBS and stored in mounting media or imaged immediately.

Antibodies used are: Cx40 (Catalog number: Cx40-A, Alpha Diagnostics, 1:500 dilution), Dach1 (Catalog number: 10194-1-AP, Proteintech, 1:500 dilution), VegfR2 (Catalog number: AF644, R&D Systems, 1:250 dilution), Endomucin (Catalog number: 14-5851-82; Thermo Fisher Scientific, 1:500 dilution) and alpha-smooth muscle actin (Catalog number: C6198, Sigma-Aldrich, 1:500 dilution). All secondary antibodies were procured from Invitrogen and used at a dilution of 1:500.

#### 3. Imaging whole brains/heads of mouse embryos

The mice heads and brains from different developmental time-points were imaged using Olympus FV-3000 upright or inverted confocal microscope by placing the sample in a 2-well chamber (Catalog number: 154461; Nunc™ Lab-Tek™ II Chamber Slide™ System; Thermo Fisher Scientific). Whole brains and heads were scanned using 4X and 10X air objectives, and multiple stacks were captured. The captured image stacks were processed in ImageJ and stitched together using Adobe Photoshop software.

The embryonic heads at e16/e16.5 were imaged using Olympus FV-3000 5L Upright microscope. The pre-fixed embryonic heads were attached to a 35mm plastic petri dish using agarose gel prior to imaging. The whole head imaging was performed using 4X air objective and watershed images were captured with 10X air objective or 20X water dipping/immersion objective and 1.5X optical zoom with brains immersed in 1X PBS.

### Visualization of pial arteries in live mice

#### 1. In vivo imaging of embryonic pial arteries

##### 1.1 Immobilization of live embryo

*Cx40CreER; Rosa26^TdTomato^*; *Tie2::eGFP::Claudin5* pregnant dams with embryos at developmental stages between embryonic day (e)16-e17 were anesthetized with a subcutaneous injection of Ketamine-Xylazine (Ketamine (80 mg/kg); Xylazine (5 mg/kg)) cocktail. The dams were then placed in a supine position on a regulated heating pad (TC1000, CWE) that was positioned on top of a 3D-printed baseplate (as shown in Figure **1A**). A small incision was placed along the midline of the abdomen. With a gentle retraction of the abdominal skin and wall, a single uterine horn within the abdominal cavity was isolated. The uterus was surgically opened and a single embryo along with the yolk sac was isolated and placed into a 35mm petri dish. The embryo still connected to the dam was immobilized by submerging the bottom part with 1% agarose, as shown in Figure **1A**. The immobilized embryo is then submerged in dissection media (DMEM/F12, catalog number: 21041025, Thermo Fisher Scientific + 10% FBS, catalog number: RM10432-500ML, HiMedia), and the media is maintained at 37°C using a custom-made Arduino controlled media heating and circulating device.

##### 1.2 Live embryonic imaging

The setup along with anesthetized dam was placed under Macro Zoom microscope MVX10-Olympus for large scale imaging and under FV3000 upright confocal Microscope-Olympus for high resolution imaging using 20X water dipping objective. In Macro Zoom Microscopy, the entire observable depth was scanned whilst obtaining videos of 1 minute spanning a duration >2 hours in a time-lapse manner. In confocal microscopy, the entire observable depth was scanned with a step size of 1 micron. A total of 4 embryos were imaged under the Macro zoom microscope and 5 embryos were imaged using confocal microscope. Out of these embryos, the artery cell dynamics was captured in 2 brains from each category.

##### 1.2 Image processing

Image stacks obtained from Macro zoom microscopy was registered using ImageJ plugin (*StackReg*) and the registered stack was de-convolved using ImageJ plugin (*DeconvolutionLab2*) and finally maximum intensity projection was obtained as shown in Figure **1D**. The image stacks obtained from the confocal microscopy were processed similarly except the deconvolution step.

The MCA and ACA growth fronts were automatically traced and the length of the traced path was measured using Simple Neurite Tracer (SNT v4.1.13). The distance of single cells from an artery was measured manually using ImageJ.

#### 2. In vivo imaging adult pial arteries

##### 2.1 Craniotomy

*Cx40CreER; Rosa26^TdTomato^*; *Tie2::eGFP::Claudin5 and VegfR2^L/L^;Cx40CreER; Rosa26^TdTomato^*; *Tie2::eGFP::Claudin5* adult mice (3-5 months) underwent craniotomy to obtain optical access to the pial layer of the brain. Mice were anesthetized under 2% Isoflurane (Abbott, North Chicago, IL, USA) and were placed on a stereotaxic mount (Item number 51547; Stoelting, Wood Dale, IL, USA) with the skull held tightly using ear bars. The mice were kept anesthetized with 2% Isoflurane using a nose-cone. 5mg/kg Dexamethasone in 1X PBS was administered subcutaneously and the mice kept warm using a heating pad (TC1000, CWE). The eyes were covered with an eye ointment and the hair above the skull removed using a hair trimmer. The surgical area was thoroughly wiped clean with 70% ethanol. Sterile surgical tools were used for the surgery. A small incision was made using a scalpel blade and the exposed skin was removed. The connective tissue covering the skull in the exposed area was scraped off using scalpel blade and Q-tips. Using a high-speed electrical drill (burr of diameter ∼0.25mm), a window of ∼4mm diameter was carefully etched on the skull. The skull within the window was eventually detached from the rest of the skull after repeated drilling and was carefully lifted using a pair of fine forceps (Catalog number: 11255-20, Dumont #55) without damaging the dura. During the process of drilling, the skull was constantly irrigated with cortex buffer (Concentration; in mM: NaCl–125, KCl–5, glucose–10, HEPES–10, CaCl_2_–2 and MgCl_2_–2. pH set to 7.35).^32^ The exposed dura was aspirated with cortex buffer until there is no bleeding and the window looked clear. An optical plug made up of two 4 mm coverslip and one 5 mm coverslip (CS-5R & CS-4R; Warner Instruments, Hamden, CT, USA) glued together (with NOA 81, Norland Optical Adhesive) was placed on the exposed dura. The optical plug was gently pressed using a toothpick such that the 5mm cover slip was touching the skull bone and then cyano-acrylic glue was applied across the perimeter of the 5mm coverslip attaching it to the skull. At this stage we had an optical window of 4mm. An aluminium head bar with circular hole of 8mm was placed onto the skull aligned to the previously made optical window. Next, dental cement (Catalog number: 56877, 3M Relyx U200) was applied over the exposed skull using a sterile toothpick while being careful to not get it on the optical window. Once the dental cement was completely dry, the animal was placed on a warm pad in a clean recovery cage. Animals undergoing craniotomy were maintained on meloxicam (2 mg/kg) and enrofloxacin (5 mg/Kg body weight) for 3-4 days until full recovery. Mice were further allowed to recover for at least 2 weeks post-surgery before the start of any experiment. Mice with an optical window can be imaged longitudinally for 2-3 months until the optical window starts to lose its clarity.

##### 2.2 Live Adult Imaging

To perform in vivo imaging, mice were anesthetized through intraperitoneal injection of Ketamine (80mg/Kg body weight) – Xylazine (5mg/Kg body weight) cocktail. The eyes were covered with an eye ointment and the mice were placed onto a custom-made baseplate and immobilized using the implanted metal head plate to this baseplate. A heating pad (37°C) was placed under the mouse to maintain the body temperature. Subsequently, the baseplate was placed on a motorized XY-table of an upright confocal microscope (FV3000 upright confocal Microscope-Olympus). The entire optical window was imaged using 10X air objective (0.4 NA) and selected pial regions with collateral arteries were imaged with 20X water dipping objective (0.5 NA). A total of N=3 adult mice per group (control and *VegfR2* artery endothelial specific knockout) were imaged longitudinally for a period of 1 month.

##### 2.3 Hypoxia regime

The *Cx40CreER; Rosa26^TdTomato^; Tie2::eGFP::Claudin5* mice (N=3) were exposed to hypoxia by gradually lowering the fraction of inspired oxygen (FiO_2_) by 1% per day over a period of 2 weeks to acclimatize them, and then maintained at 7% FiO_2_ for an additional 2 weeks (Figure **S12C**). The hypoxia chamber (Biospherix Ltd.) was continuously supplied with nitrogen gas as needed to maintain the desired oxygen levels, and CO_2_ levels were kept below 600 ppm throughout the experiment by changing the number of open holes (0.7 cm diameter) provided on each of the walls. Oxygen and CO_2_ levels were constantly monitored using inbuilt sensors (Biospherix; ProOx Model P360 and ProCO2 Model P120ppm).

##### 2.3 Image processing

The 10X image stacks obtained from the confocal microscopy of the cranial window were registered using ImageJ plugin (*StackReg*) and maximum intensity projection was obtained. The region of interest from the 20X image stacks were cropped out slice by slice and stitched in Adobe Photoshop to avoid the overlaying microvascular network in order to focus on the pial collaterals.

## QUANTIFICATION AND STATISTICAL ANALYSIS

### Number of collateral arteries per hemisphere

In *Cx40CreER^+^* (TdTomato) labelled brains, the collateral number was quantified based on the number of contiguous TdTomato^+^ vessels (Td-contiguous) connecting MCA and ACA. Data shown in Figure **4K** was summarized from 2 control and 3 artery-specific *VegfR2* knockout. Data shown in Figure **S2H** was summarized from e13.5 (N=7), e14.5 (N=9), e15.5 (N=6), e18.5 (N=6) and P0 (N=5) embryos and neonates. Data shown in Figure **S3H** was summarized from e13.5 (N=3), e14.5 (N=3), e15.5 (N=2), e18.5 (N=2) and P0 (N=5) embryos and neonates. Data shown in Figure **S9E** is quantified by counting number of Cx40 immunostained collateral arteries between MCA and ACA, and is summarized from N=4 control and N=7 capillary specific *Dach1* knockouts. An unpaired *t* test was done to test significance statistically. Data were plotted using GraphPad Prism and presented as mean ± standard deviation. The p values in statistical analyses have been described as follow: ns (not significant), p > 0.05; *p < 0.05; **p < 0.01; ***p < 0.001; and ****p < 0.0001.

### Microvascular tracks and their coverage with TdTomato^+^ arteries per watershed

To analyse the Microvascular (MV) tracks, the watershed regions of the e16 (9 watershed from N=9 embryos) and e16.5 (7 watershed from N=5 embryos) brains were imaged using 10X air objective at 1.5X optical zoom. The region of interest (ROI) of comparable width (600-800 microns) within the watershed was taken for further analyses. The eGFP^+^ MV tracks were traced using Simple Neurite Tracer (SNT v4.1.13). The total MV length and area of the ROI were measured using SNT and ImageJ respectively, to calculate the total MV length per unit area of the watershed. Further, to measure TdTomato coverage of the MV tracks, the TdTomato^-^ vessel segments on the MV track were traced and their lengths measured using SNT to quantify the fraction of TdTomato coverage. Both, the MV tracks and TdTomato coverage were quantified by going through multiple z-slices and data were presented as mean ± standard deviation. Two-tailed unpaired *t* test was done to test statistical significance using GraphPad Prism software (p=0.4085 for %TdTomato coverage in the MV tracks in Figure 2O and p=0.009 for normalized total MV length per watershed in Figure 2J). The p values in statistical analyses have been described as follow: ns (not significant), p > 0.05; *p < 0.05; **p < 0.01; ***p < 0.001; and ****p < 0.0001.

### Lineage contribution towards pial collaterals during embryogenesis

Cx40 immunostained pial collateral segments from either *Cx40CreER; Rosa26^TdTomato^* or *ApjCreER; Rosa26^TdTomato^* brains were imaged at 10X with 2.12 optic zoom. Regions of interest were manually drawn to mark the collateral artery segments. Lineage contribution was determined by quantifying TdTomato^+^ coverage on the collateral segment (Figure **3**). This was performed by thresholding the images with Otsu algorithm in ImageJ. The area fraction of the TdTomato^+^ area per collateral segment was calculated using the Measure tool in ImageJ. Contribution of the artery and capillary derived ECs in collaterals was quantified by calculating the percentage of collateral area covered by TdTomato^+^ signal. 45 collateral segments (of an approximate length of 600-800µm) from 7 *Cx40CreER*-lineage traced brains (biological replicate) and 57 collateral segments from 8 *ApjCreER-*lineage traced brains were analysed. The data were presented as mean ± standard deviation using GraphPad Prism.

### Pial artery coverage in whole brains and in the collateral zone

The image stacks were obtained using confocal imaging of whole embryonic brains with 10X objective, stitched using ImageJ plugin (*Stitching*) and max intensity projection (MIP) was obtained. The MIPs were used for further analysis wherein one complete hemisphere was cropped out and aligned vertically to the major axis of its own elliptical boundary. Using a custom ImageJ macro, a line of the length of the major axis was scanned across the width of the image and intensity profile for every pixel along the horizontal axis was generated. Total number of local maxima (peaks) for each intensity profile was obtained using the built-in ImageJ macro function *Array.findMaxima (array, tolerance). The* tolerance used was 2 times the average background intensity. The number of peaks thus obtained was plotted across the width and the plot was fitted to a Gaussian function using OriginPro 2024b software. The fitted curve was used as a representative of the arterial coverage per hemisphere. The fit curves obtained from various mice were aligned based on the Gaussian peaks and averaged to get the representative curves for various experimental groups. We considered a region of 1mm (-0.5mm to +0.5 mm) across the Gaussian peak as the collateral zone (watershed). The area under the curve within this region was used as a measure for the extent of the collateral network. 9 hemispheres from 9 embryos (e18.5) were analysed per group (control and artery-specific knockout for *VegfR2*). To test the statistical significance of the differences observed in pial artery coverage (Figure **4E**), a Kolmogorov-Smirnov (KS) test was done. D value= 0.569, p= 2.2066 X 10^-141^. To test the statistical significance of the area under the curve (pial collateral artery network shown in Figure **4F**), two-tailed unpaired *t* test was done (p=0.0001) using GraphPad Prism. Similar analyses were performed in Figures **S7** and **S10**. Two-tailed unpaired *t* test was performed using the same. In all cases, data were presented as mean ± standard deviation and each dot in Figures **4F****, S10G, S10K** and **S10O** represents a brain (biological replicate). The p values in statistical analyses have been described as follow: ns (not significant), p > 0.05; *p < 0.05; **p < 0.01; ***p < 0.001; and ****p < 0.0001.

### Efficiency of TdTomato recombination

Segments of collateral arteries, proximal arteries (PA) and distal arteries (DA) from the image stacks taken with 20X objective were obtained from live adult imaging of *Cx40CreER*; *Rosa26^TdTomato^*; *Tie2::eGFP::Claudin5* mice. TdTomato^+^ and TdTomato^-^ cells were counted in each artery segment (Figure **5D**) using ImageJ. Each segment was of an approximate length of 600-800µm.The recombination efficiency was measured as ratio of TdTomato^+^ cells and total number of cells in the segment. A total of 10 collateral segments, 10 PA segments and 10 DA segments from 4 mice were analysed to obtain the recombination efficiency of TdTomato. One-way ANOVA was done to test significance statistically (Tukey’s mean comparison test, PA vs. collateral, p<0.0001; DA vs. collateral, p<0.0001; PA vs. DA, p=0.08). GraphPad Prism was used to visualize the data as mean ± standard deviation. The p values in statistical analyses have been described as follow: ns (not significant), p > 0.05; *p < 0.05; **p < 0.01; ***p < 0.001; and ****p < 0.0001.

### Impact of arterial VegfR2 deletion on collateral formation

To assess the effect of arterial VegfR2 deletion on pial collateral development, *VegfR2^L/L^; Cx40CreER*; *Rosa26^TdTomato^*; *Tie2::eGFP::Claudin5* mouse strain was used. The watershed regions of e16.5 brains from control (7 watersheds from N=5 embryos) and knockout (5 watersheds from N=5 embryos) were imaged using 10X air objective at 1.5X optical zoom. Microvascular tracks and their coverage with TdTomato^+^ arterial cells were quantified using ImageJ (Figures **4K** and **S8A-S8C**) following above-mentioned method. Capillary density was measured by thresholding the images and obtaining the area fraction using ImageJ (Figures **S8D-S8E**). Each dot in Figure S8E represents brain (biological replicate). For all cases, data were plotted as mean ± standard deviation and two-tailed unpaired t test was done to test the statistical significance using GraphPad Prism software (p=0.0004 for normalized total MV length per watershed, p=0.0004 for %TdTomato coverage, and p=0.73 for capillary density). The p values in statistical analyses have been described as follow: ns (not significant), p > 0.05; *p < 0.05; **p < 0.01; ***p < 0.001; and ****p < 0.0001.

### Quantification for deletion of *Dach1*

*Dach1* deletion from the capillaries was validated in brains at embryonic day 18.5 by immunostaining with antibodies for Dach1, VegfR2, and SMA along with nuclear marker DAPI (Figure **S9C**). Dach1 expression was quantified manually using ImageJ by calculating corrected total cell fluorescence (CTCF) from individual DAPI^+^ capillary nucleus. 5 regions of interest (ROI) from each of the control (N=2) and 4 ROIs from each knockout brains (N=3 biological replicates) were used for analyses. The data were plotted as mean ± standard deviation and two-tailed unpaired *t* test was done to test the statistical significance using GraphPad Prism (p<0.0001). The p values in statistical analyses have been described as follow: ns (not significant), p > 0.05; *p < 0.05; **p < 0.01; ***p < 0.001; and ****p < 0.0001.

## AUTHORS CONTRIBUTION

SD conceived the idea and designed the experiments. SK performed intravital imaging of embryos and adults. SK, SG, NS, VS, MD performed embryonic experiments. SD, SK, SG, NS wrote the manuscript. All authors contributed to editing the manuscript.

## ACKNOWLEDGEMENTS

We thank the following facilities for their technical support: Animal Care and Resource Center at NCBS, Mouse Genome Engineering Facility at NCBS, Central Imaging and Flow Cytometry Facility at NCBS. We sincerely thank Dr. Dritan Agalliu (Columbia University) and Dr. Kristy Red-Horse (Stanford University) for generously sharing *Tie2::eGFP::Claudin5* and *Dach1^fl/fl^* mice respectively.

## Supplementary figures and figure legends

**Supplemental Figure 1:**
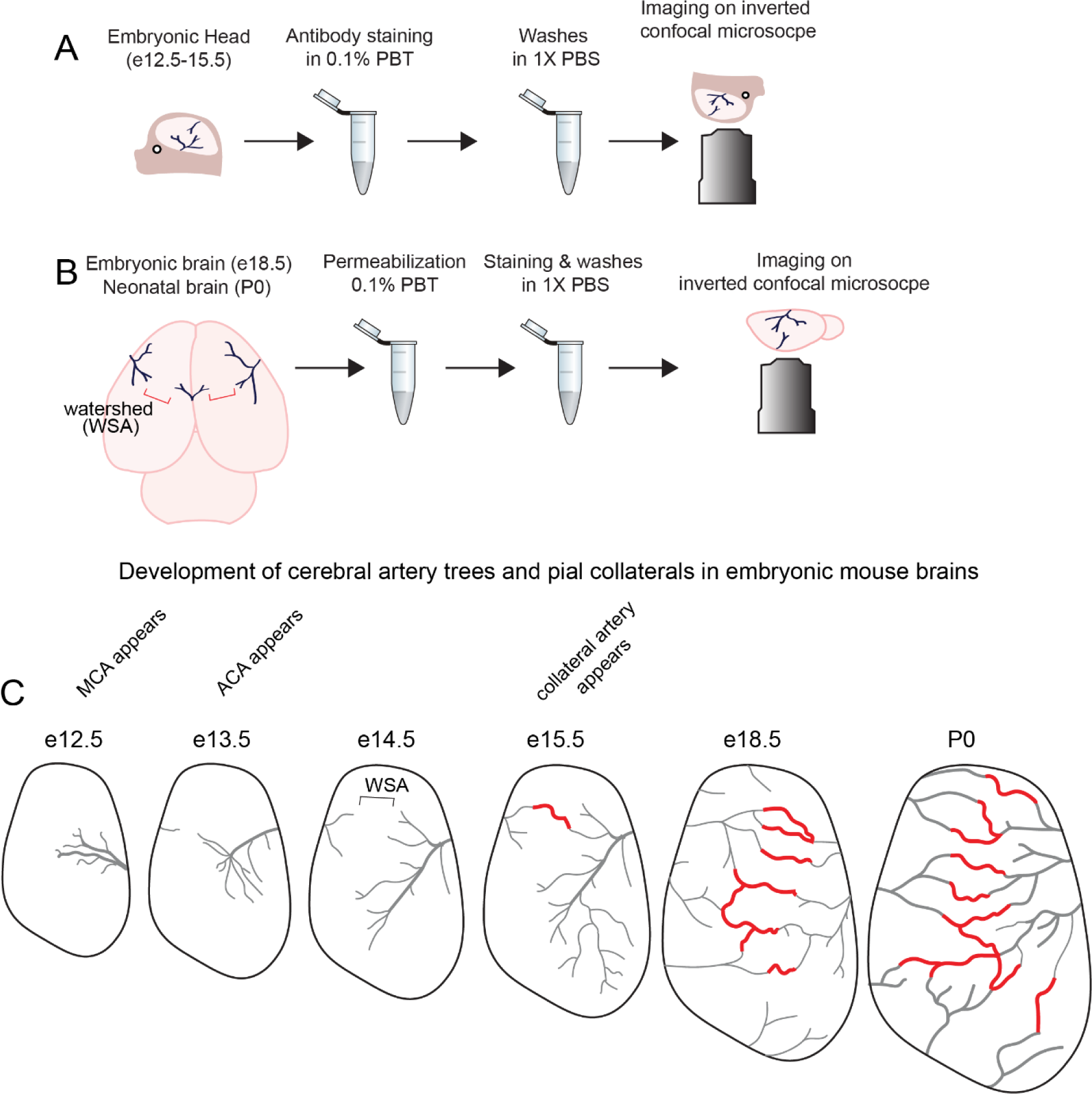
Visualization of pial arteries and collateral arteries, related to Figure 1. Schematic summarizes key steps of immunostaining methods used to image pial artery network in (**A**) mouse embryonic heads and (**B**) mouse embryonic/neonatal brains with confocal microscope. (**C**) Schematic shows findings from **A** and **B** in one hemisphere. Grey represents cerebral arteries, Red represents collateral artery connections between cerebral arteries. Brackets show watershed area (WSA), MCA appears at e12.5, ACA appears at e13.5, and the foremost pial collateral arteries appear at e15.5. These pial collaterals then increase in number till e18.5/P0. e, embryonic day; P, postnatal day; MCA, middle cerebral artery; ACA, anterior cerebral artery.

**Supplemental Figure 2:**
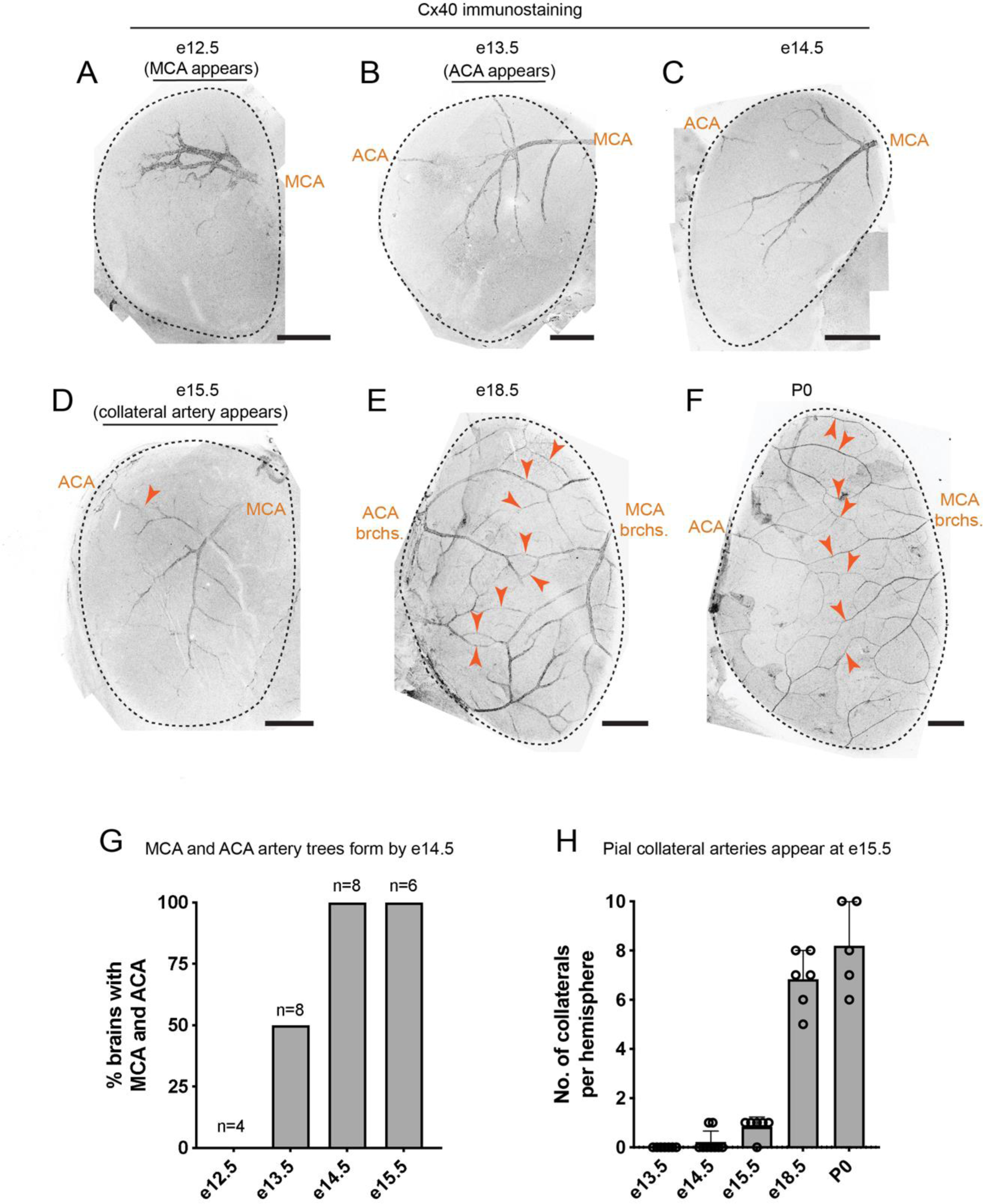
Detection of pial collaterals with whole brain immunostaining, related to Figure 1. (**A**-**F**) Representative images of right hemispheres from developing mouse embryos immunostained for Connexin-40 (Cx40). Cx40 marks all artery endothelial cells. MCA appears at e12.5, ACA appears at e13.5 and collaterals between MCA and ACA appear at e15.5. Arrowheads point to collateral arteries between MCA and ACA. (**G**) Quantification of Cx40 immunostained brains from different developmental stages reveals that both MCA and ACA have formed by e14.5. (**H**) Quantification of Cx40^+^ collateral arteries per hemisphere shows that the first pial collaterals appear at e15.5. N=7 (e13.5), 9 (e14.5), 6 (e15.5), 6 (e18.5) and 5 (P0 brain). Each dot represents an individual brain. e, embryonic day; P, postnatal day; MCA, middle cerebral artery; ACA, anterior cerebral artery; brch., branch. Scale bars: 500µm

**Supplemental Figure 3:**
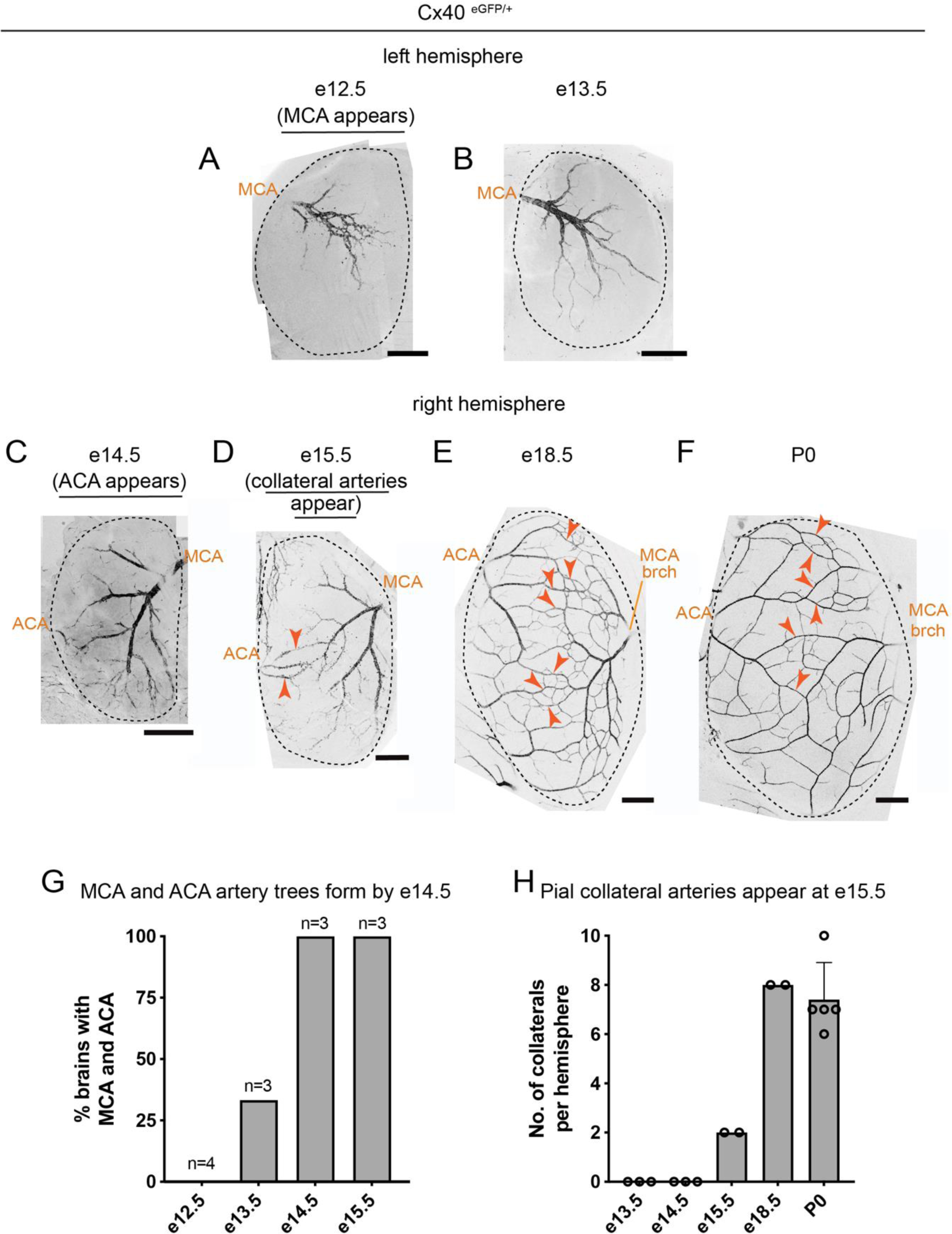
Detection of pial collaterals with *Cx40* reporter activity, related to Figure 1. (**A**-**F**) Representative images of brains from developing *Cx40^eGFP/+^* mouse embryos. Connexin-40 (Cx40) marks all artery endothelial cells. MCA appears at e12.5, ACA appears at e14.5 and collaterals between MCA and ACA appear at e15.5. Arrowheads point to collateral arteries between MCA and ACA. (**G**) Quantification of *Cx40^eGFP/+^* brains from different developmental stages reveals that both MCA and ACA have formed by e14.5. (**H**) Quantification of eGFP^+^ collateral arteries per hemisphere shows that the first pial collaterals appear at e15.5. N=3 (e13.5), 3 (e14.5), 2 (e15.5), 2 (e18.5) and 5 (P0 brain). Each dot represents an individual brain. e, embryonic day; P, postnatal day; MCA, middle cerebral artery; ACA, anterior cerebral artery; brch., branch. Scale bars: 500µm

**Supplemental Figure 4:**
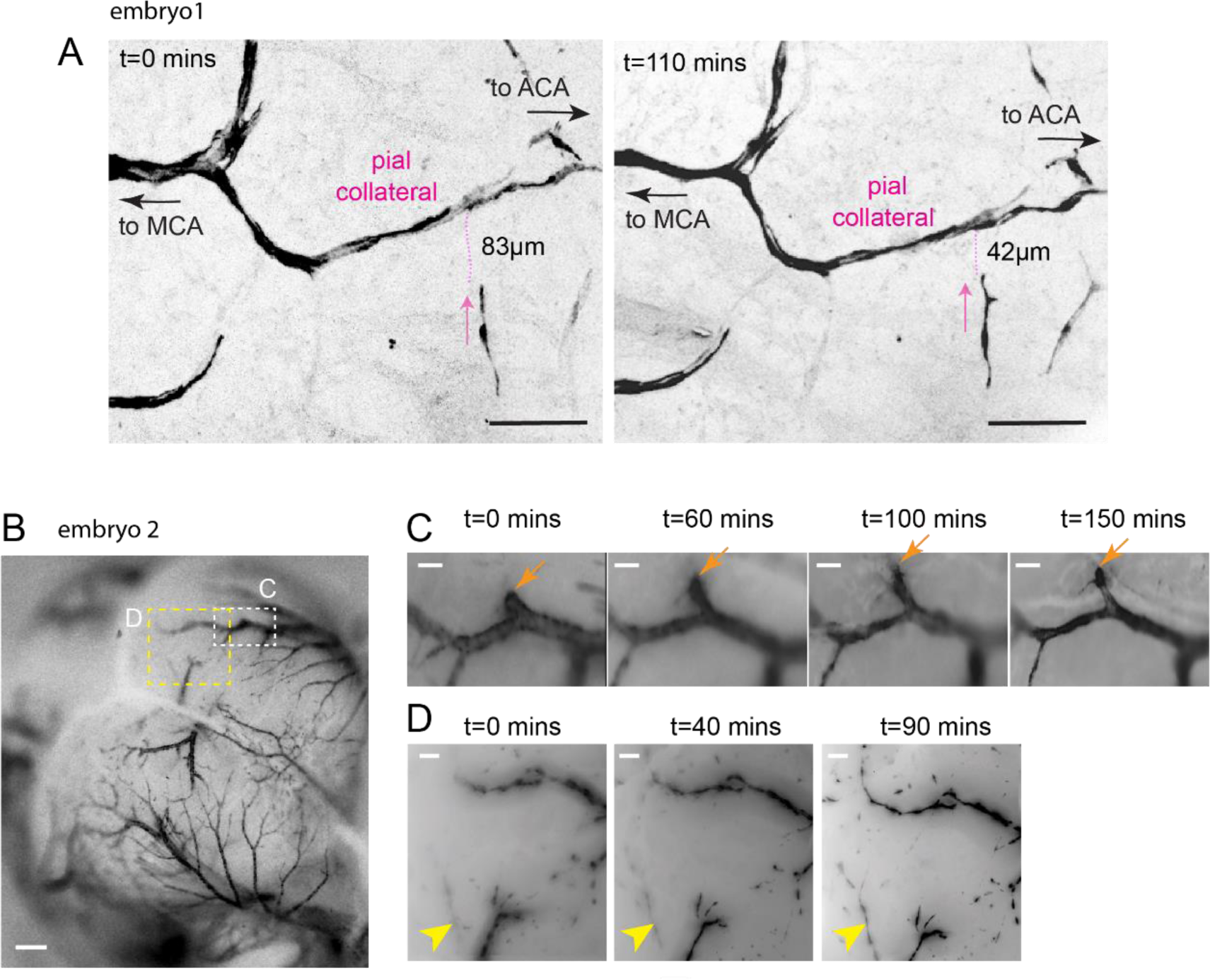
Visualization of pial artery cell dynamics on microvascular tracks, related to Figure 1. (**A**) Confocal image of e16 embryonic collateral zone showing single recombined artery endothelial cell migration towards the pial collateral within 110mins. Dotted line is the length of micro-vessel between the single recombined cell and the pial collateral. Arrow shows direction of single recombined artery cell migration. (B) Stereo-image of a whole e16 brain showing both hemispheres and the pial watersheds. Insets C and D show artery growth fronts of MCA and ACA respectively. (C) Time-lapse stereo images show growth of MCA branch within 150 minutes. (D) Time-lapse stereo images show appearance of an ACA branch within 90 minutes. MCA, middle cerebral artery; ACA, anterior cerebral artery. Tamoxifen was injected at e13.5. Scale bars: **A**, **B** 100μm; **C**, **D**, 50μm

**Supplemental Figure 5:**
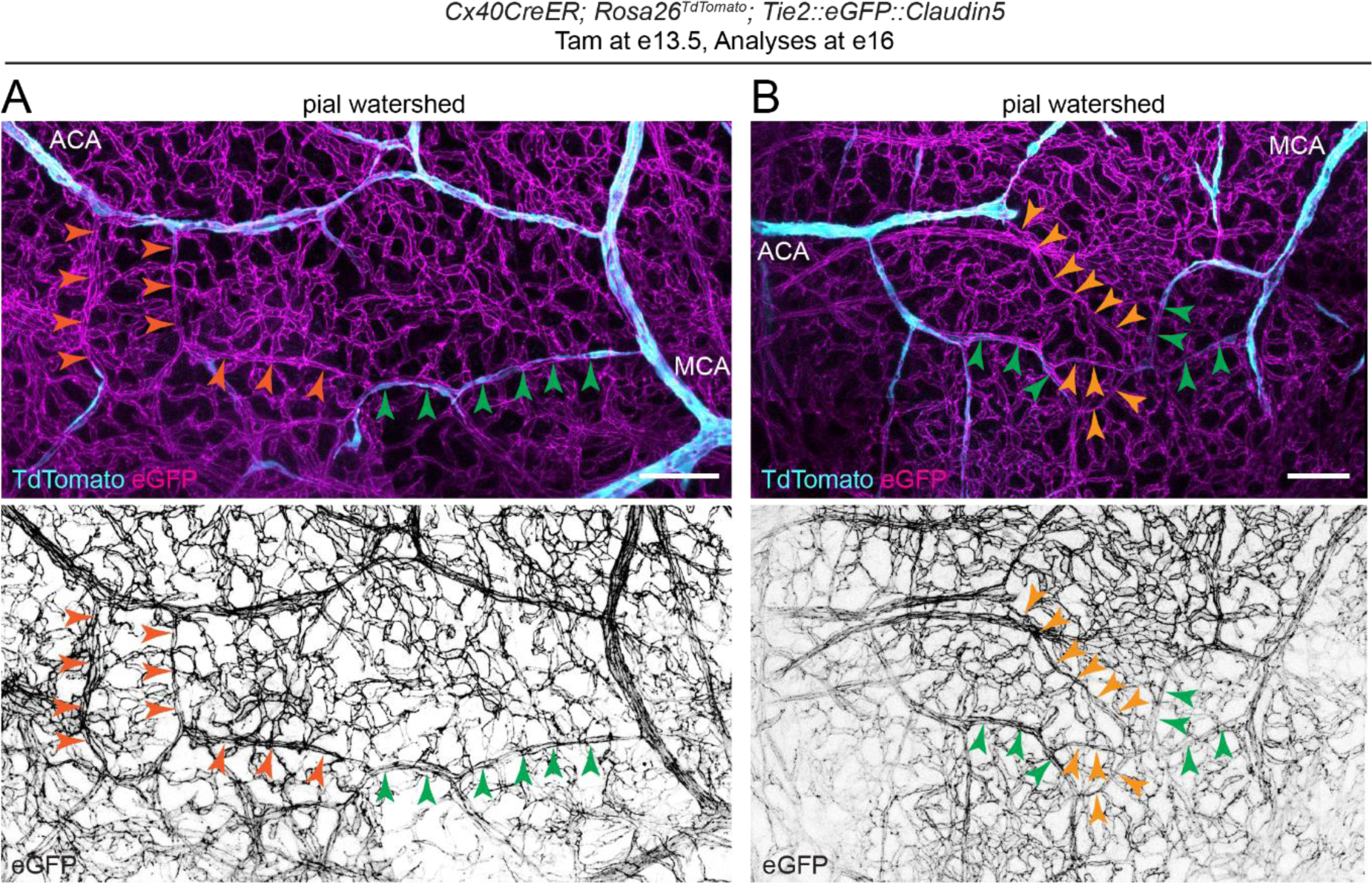
Analyses of MV tracks in pial watershed, related to Figure 2. (**A**, **B**) Confocal images of pial watershed area from *Cx40CreER*; *Rosa26^TdTomato^*; *Tie2::eGFP::Claudin5* brains, show TdTomato^+^ arteries in cyan and eGFP^+^ microvasculature in magenta. Microvascular (MV) tracks are shown using arrowheads. Orange arrowheads point to MV tracks not covered by artery cells. Green arrowheads point to MV tracks partially or completely covered by artery cells. Tamoxifen was administered at e13.5, and brains were analysed at e16. Scale bars: 100µm

**Supplemental Figure 6:**
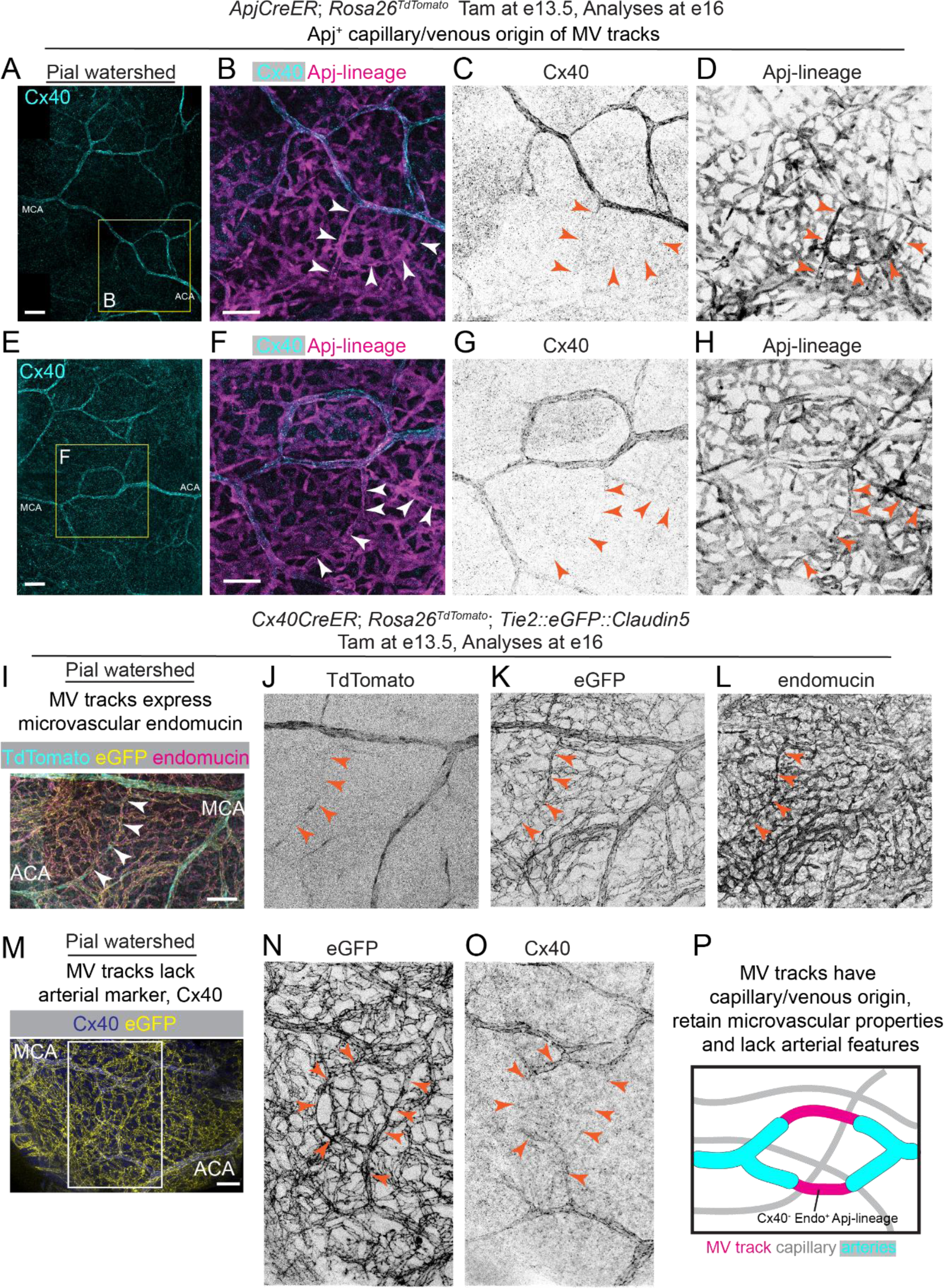
Molecular characterization of MV track during pial collateral formation, related to Figure 2. (**A**-**H**) Confocal images of pial watershed from *ApjCreER; Rosa26^TdTomato^* embryos. MV tracks (arrowheads) lack expression of artery endothelial cell marker Cx40 and show expression of TdTomato, an indicator for *Apj*-lineage. (**I**-**L**) Confocal images show expression of Endomucin protein, a microvascular marker, on TdTomato^-^ eGFP^+^ MV tracks. Arrowheads point to MV tracks. (**M**-**O**) Confocal images of pial watershed from *Cx40CreER*; *Rosa26^TdTomato^*; *Tie2::eGFP::Claudin5* brains show lack of expression of Cx40 protein, an artery EC marker, on TdTomato^-^ eGFP^+^ MV tracks. Arrowheads point to MV tracks. (**P**) A cartoon of pial watershed with Endomucin^+^Cx40^-^ MV tracks. Cyan represents arteries, magenta represents MV tracks. For all experiments, Tamoxifen was administered at e13.5, and brains were analysed at e16. Scale bars: 100µm

**Supplemental Figure 7:**
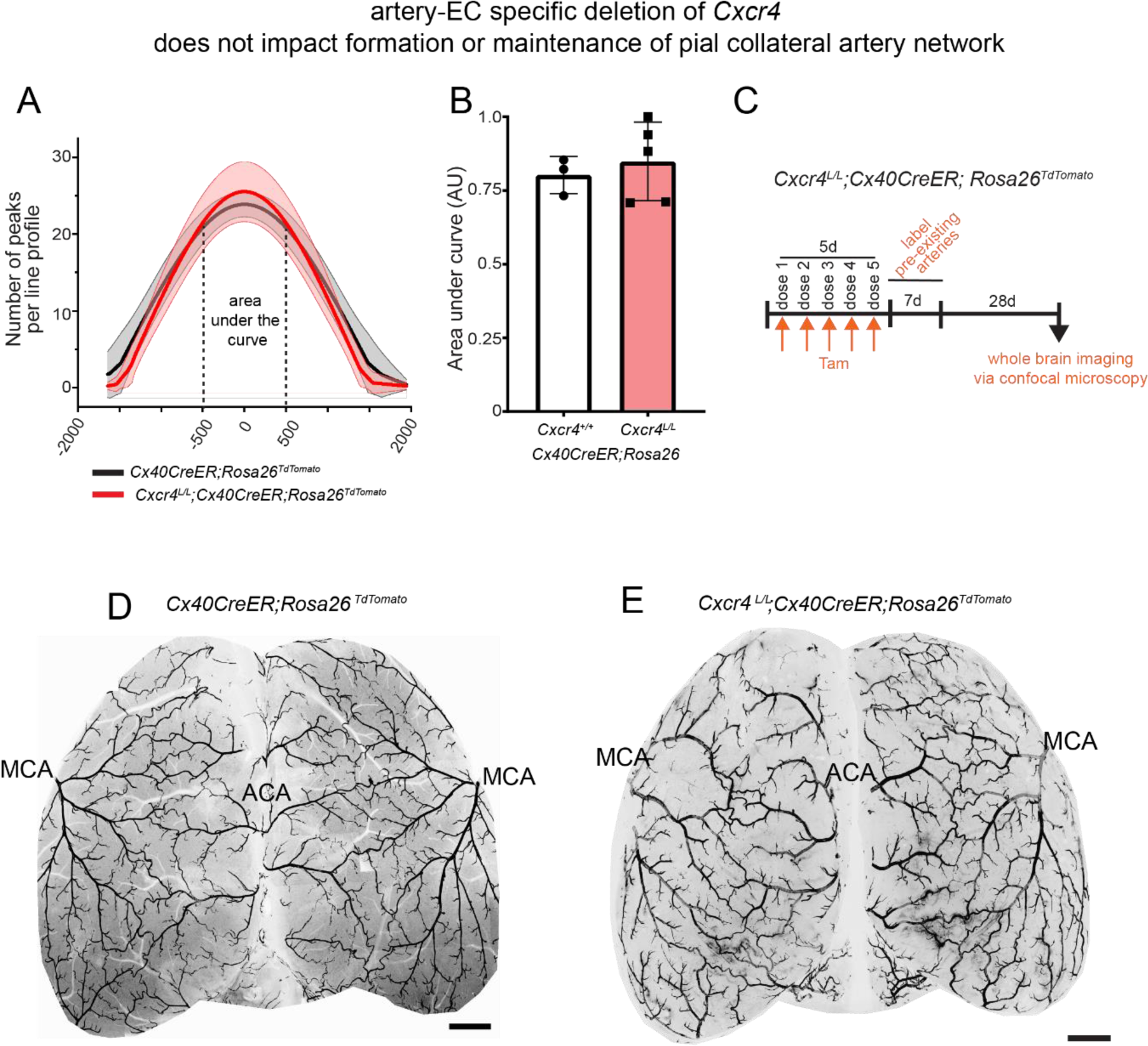
Pial artery coverage does not reduce without arterial *Cxcr4*, related to Figure 4. (**A**) Quantification of overall pial artery coverage with and without arterial *Cxcr4*. KS test performed on the distributions generated D value= 0.1944, p value= 0.0001. (**B**) Quantification of pial collateral artery network with and without arterial *Cxcr4* (N=3 for control and N=5 for knockout). Each dot represents a biological replicate. Data shown as mean ± standard deviation. Two-tailed unpaired *t* test, p=0.606. (**C**) Experimental set up to assess the effect of arterial *Cxcr4* deletion on pial artery coverage in adults. (**D**, **E**) Representative confocal images of whole adult brains (from N=5 for each genotype), genetically lineage traced for pre-existing artery endothelial cells, (**D**) with or (**E**) without arterial *Cxcr4*. ACA, anterior cerebral artery; MCA, middle cerebral artery; Tam, Tamoxifen. Scale bars: 1mm

**Supplemental Figure 8:**
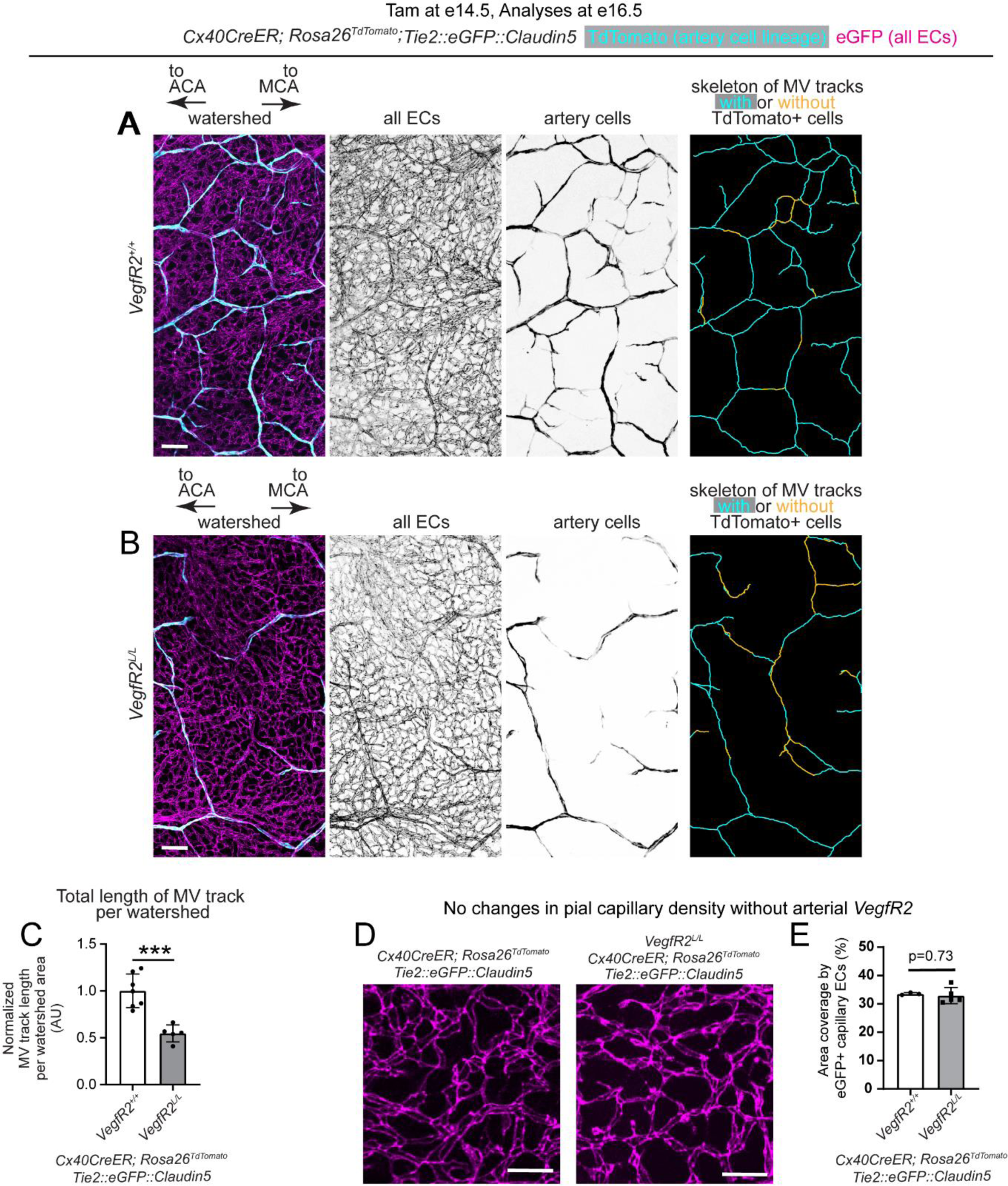
Analyses of MV tracks in pial watershed, without arterial VegfR2, related to Figure 4. (**A**, **B**) Higher magnification confocal images of e16.5 watershed region between ACA and MCA (**A**) with or (**B**) without arterial *VegfR2*. All endothelial cells are in magenta and artery endothelial cells in cyan. Tamoxifen was injected at e14.5. Skeleton of all the eGFP^+^ microvascular tracks traced in the e16.5 watershed through multiple z-slices, using Simple Neurite Tracer. Traced MV tracks occupied by TdTomato^+^ artery endothelial cells are shown in cyan and MV tracks not occupied with TdTomato is shown in yellow. (**C**) Quantification of total length of microvascular tracks per watershed area at e16.5 with or without arterial *VegfR2* (control: 7 watersheds from N=5 embryos, artery endothelial cell specific knockout for *VegfR2*: 5 watersheds from N=5 embryos). Two-tailed unpaired *t* test, p=0.0004. Each dot represents a selected watershed region from embryonic brain. Data shown as mean ± standard deviation. (**D**) Confocal images of eGFP^+^ capillary ECs (magenta) in e16.5 pial watershed with or without arterial *VegfR2*. (**E**) Quantification of capillary EC density with (N=3 brains) or without (N=5 brains) arterial *VegfR2*. Two-tailed unpaired *t* test, p=0.73. Each dot represents a biological replicate. Data shown as mean ± standard deviation. ***p < 0.001. Tam, Tamoxifen; MCA, middle cerebral artery; ACA, anterior cerebral artery; MV, Microvascular; e, embryonic day; EC, endothelial cell; AU, Arbitrary units. Scale bars: **A**, **B**, 100 µm; **D**, 50µm.

**Supplemental Figure 9:**
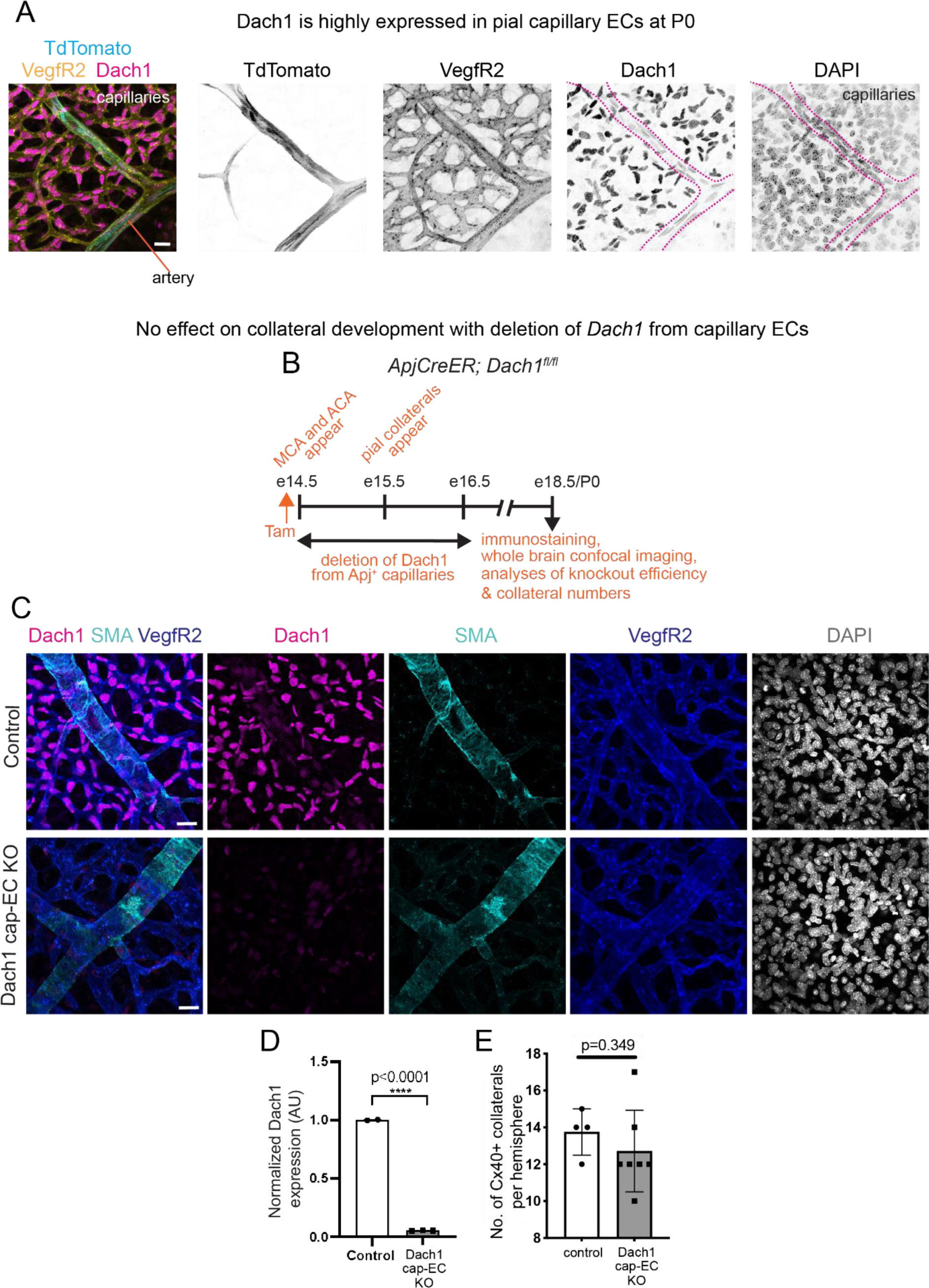
Dach1-independent pial collateral development, related to Figure 4. (**A**) Dach1 expression (green) in capillaries and arteries in P0 brains. Dotted line marks the TdTomato^+^ (cyan) arterial boundary. VegfR2 marks all vasculature. DAPI marks nuclei. (**B**) Experimental design showing Tamoxifen injection at e14.5, and analyses of collateral numbers in e18.5/P0 brains. (**C**) Confocal images of pial vasculature show absence of Dach1 from capillary endothelial cells. (**D**) Quantification of *Dach1* deletion from pial capillary endothelial cells. N=2 for control and N=3 for knockout. Two-tailed unpaired *t* test, p < 0.0001. (**E**) Quantification of Cx40^+^ pial collaterals between MCA and ACA in control (*Dach1^fl/fl^*, N=4) and *ApjCreER*; *Dach1^fl/fl^* (Dach1 cap-EC KO, N=7) brains. Two-tailed Welch’s *t* test, p=0.349. Each dot in **D** and **E** represents a biological replicate. Data shown as mean ± standard deviation. ****p < 0.0001. e, embryonic day; cap-EC, capillary endothelial cell; KO, knockout; Tam, Tamoxifen. Scale bar: A, 20µm; C, 25µm

**Supplemental Figure 10:**
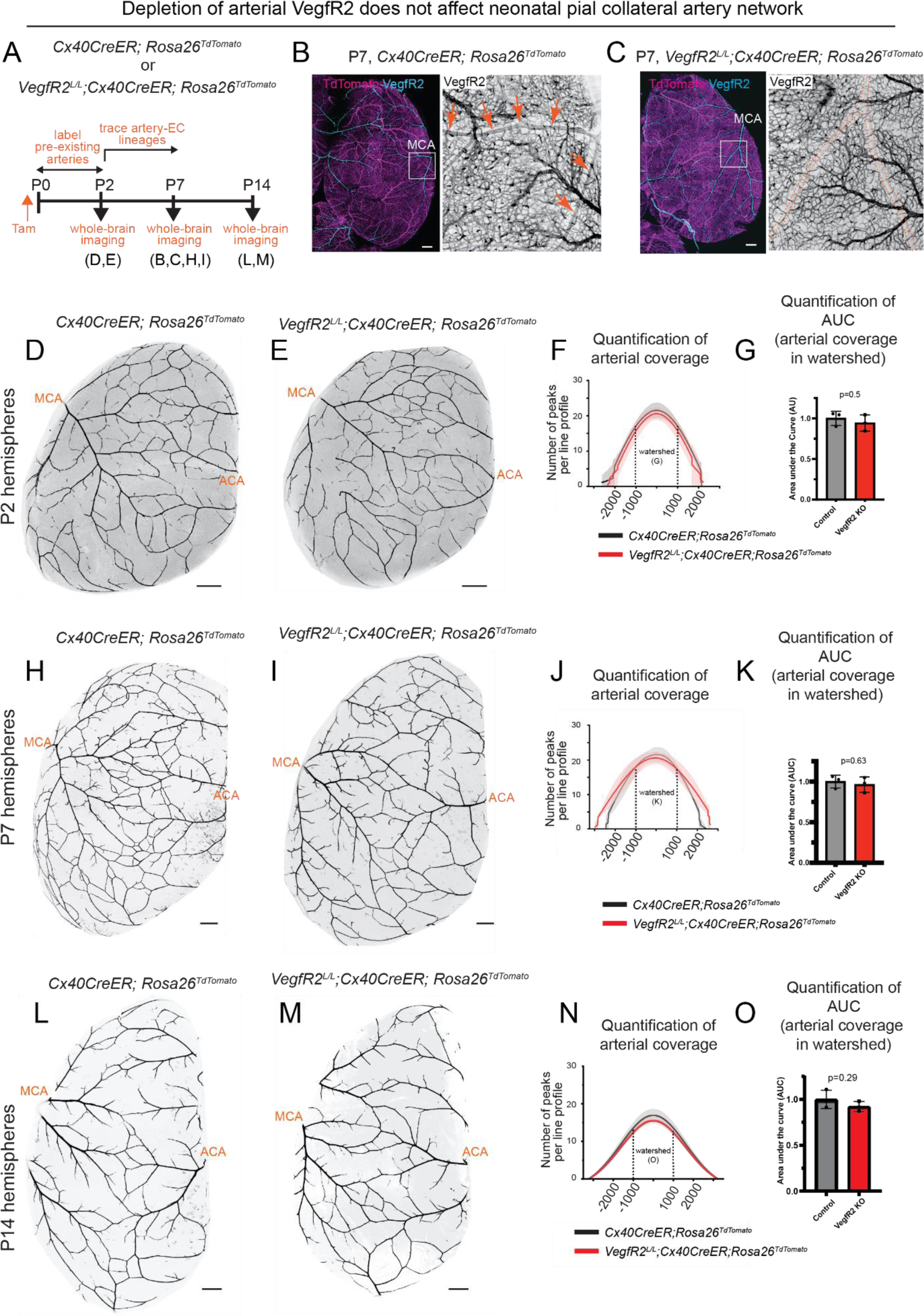
VegfR2-independent pruning of neonatal pial collaterals, related to Figure 5. (**A**) Experimental design. (**B**) Confocal imaging of whole brains immunostained for VegfR2 shows presence of the protein in pial arteries. Boxed region in left image is magnified as the right image. Arrows point to VegfR2 protein expression in MCA branches. (**C**) Confocal imaging of whole brains immunostained for VegfR2 shows absence of the protein in pial arteries (enclosed in dotted lines). Boxed region in left image is magnified as the right image. (**D**, **E**) Confocal whole P2 brain images showing TdTomato^+^ arteries (**D**) with or (**E**) without *VegfR2*. (**F**, **G**) shows quantification of (**F**) pial artery coverage across the entire P2 hemisphere or (**G**) collateral artery coverage in the watershed, in brains with or without arterial VegfR2. Two-tailed unpaired *t* test was performed with p= 0.5. (**H**, **I**) Confocal whole P7 brain images showing TdTomato^+^ arteries (**H**) with or (**I**) without *VegfR2*. (**J, K**) shows quantification of (**J**) pial artery coverage across the entire P7 hemisphere or (**K**) collateral artery coverage in the watershed, in brains with or without arterial *VegfR2*. Two-tailed unpaired *t* test was performed with p= 0.63. (**L**, **M**) Confocal whole P14 brain images showing TdTomato^+^ arteries (**L**) with or (**M**) without *VegfR2*. (**N**, **O**) shows quantification of (**N**) pial artery coverage across the entire P14 hemisphere or (**O**) collateral artery coverage in the watershed, in brains with or without arterial *VegfR2*. For all the plots, each dot (N=3 for each group) represents individual mouse brain. Two-tailed unpaired *t* test was performed with p= 0.29. P, postnatal day; KO, knockout; Tam, Tamoxifen. Scale bars: 500μm

**Supplementary Figure 11:**
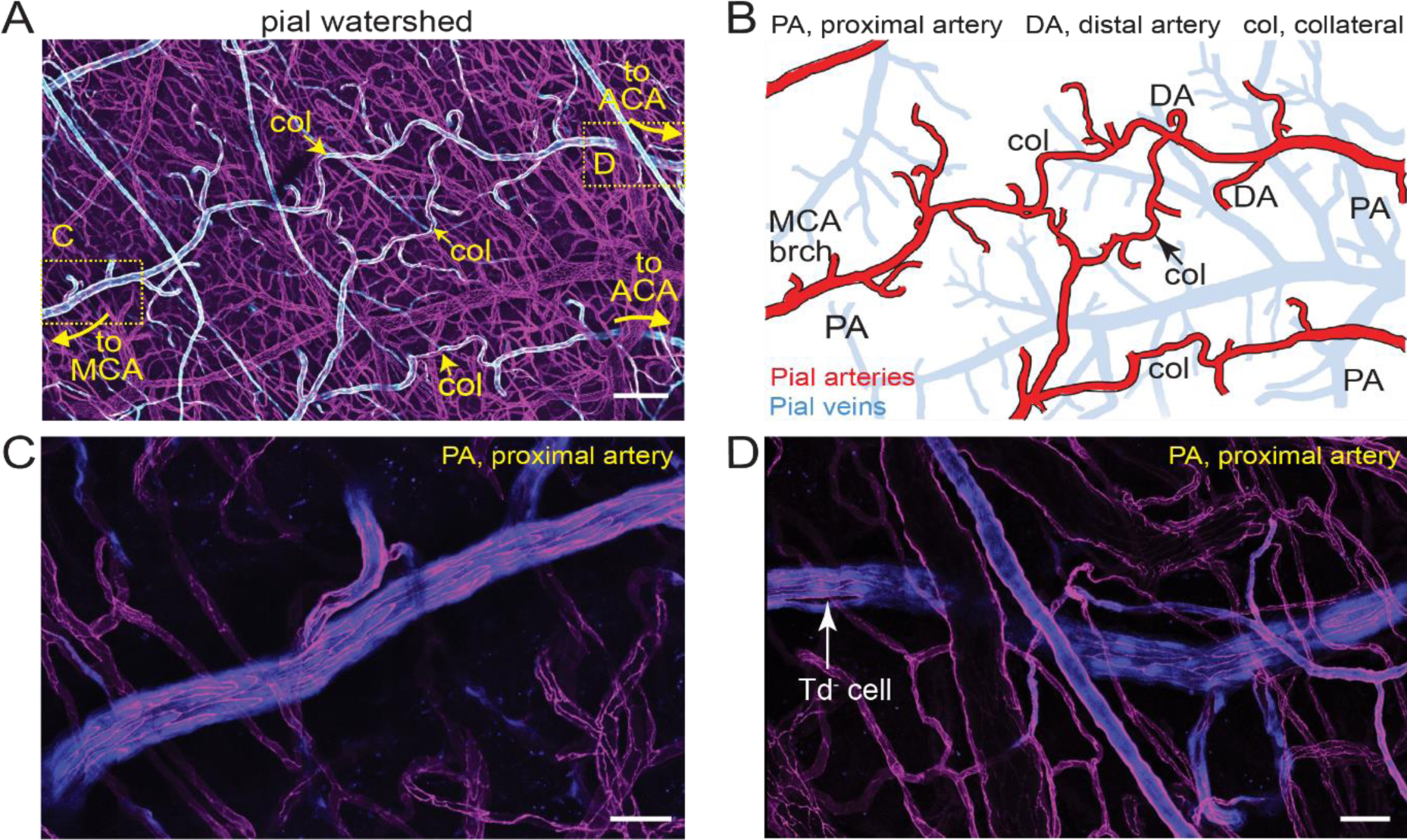
*In vivo* imaging of adult proximal arteries, related to Figure 5. (**A**-**D**) Images of pial watershed with pre-existing artery endothelial cells labelled in TdTomato (cyan) and all endothelial cells labelled in eGFP (magenta). (**A**) Confocal image from optical window created for longitudinal imaging of *Cx40CreER; Rosa26^TdTomato^; Tie2::eGFP::Claudin5* mouse as shown in Figure 5A. The image shows both pial and dural vessels. (**B**) Pial layer segmented out from **A**. red and blue show pial arteries and veins respectively. (**C**, **D**) Confocal images of pial proximal arteries from boxed regions shown in **A**. **C** and **D** are proximal arteries connected to MCA and ACA respectively. Arrow shows a TdTomato negative cell. MCA, middle cerebral artery; ACA, anterior cerebral artery; PA, proximal artery; DA, distal artery; col, collateral; brch., branch. Scale bar: A, 200µm; C, D, 50µm

**Supplemental Figure 12:**
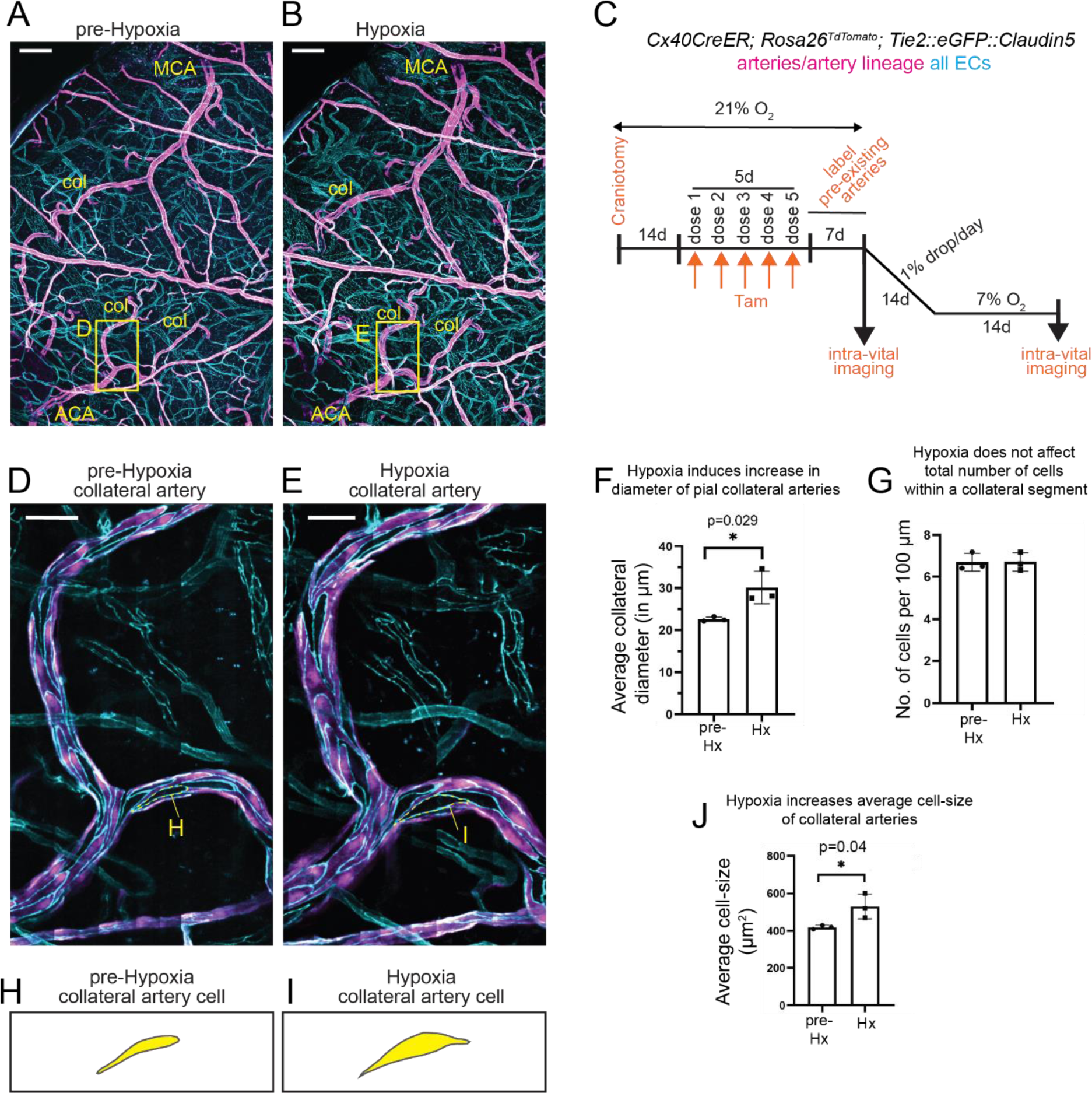
Hypoxia induces increase in size of pial collateral cells, related to Figure 5. (**A, B**) Cranial windows showing pial and dural vessels from mice (**A**) under normoxia and (**B**) hypoxia. (**C**) Experimental design using *Cx40CreER; Rosa26^TdTomato^; Tie2::eGFP::Claudin5*. (**D**, **E**) Insets shown in **A** and **B** respectively. Pial collateral artery segments from (**D**) normoxic and (**E**) hypoxic conditions. (**F**) Quantification of diameter of pial collateral arteries in before and during hypoxia. Two-tailed unpaired *t* test, p=0.029. (**G**) Quantification of total number of cells within a pial collateral artery before and during hypoxia. Two-tailed unpaired *t* test, p=0.99. (**J**) Quantification of average cell size before and during hypoxia. Two-tailed unpaired *t* test, p=0.046. (**H**, **I**) Cells marked in **D**, and **E** respectively. A representative cell from normoxic and hypoxic pial collateral segment. In **F**, **G**, and **J**, each dot represents a biological replicate (N=3). Data shown as mean ± standard deviation. *p < 0.05. MCA, middle cerebral artery; ACA, anterior cerebral artery; col, collateral artery; Hx, Hypoxia; d, day; O_2_, oxygen. Scale bars: **A**, **B**, 200µm; **D**, **E**, 50µm.

**Supplemental Figure 13:**
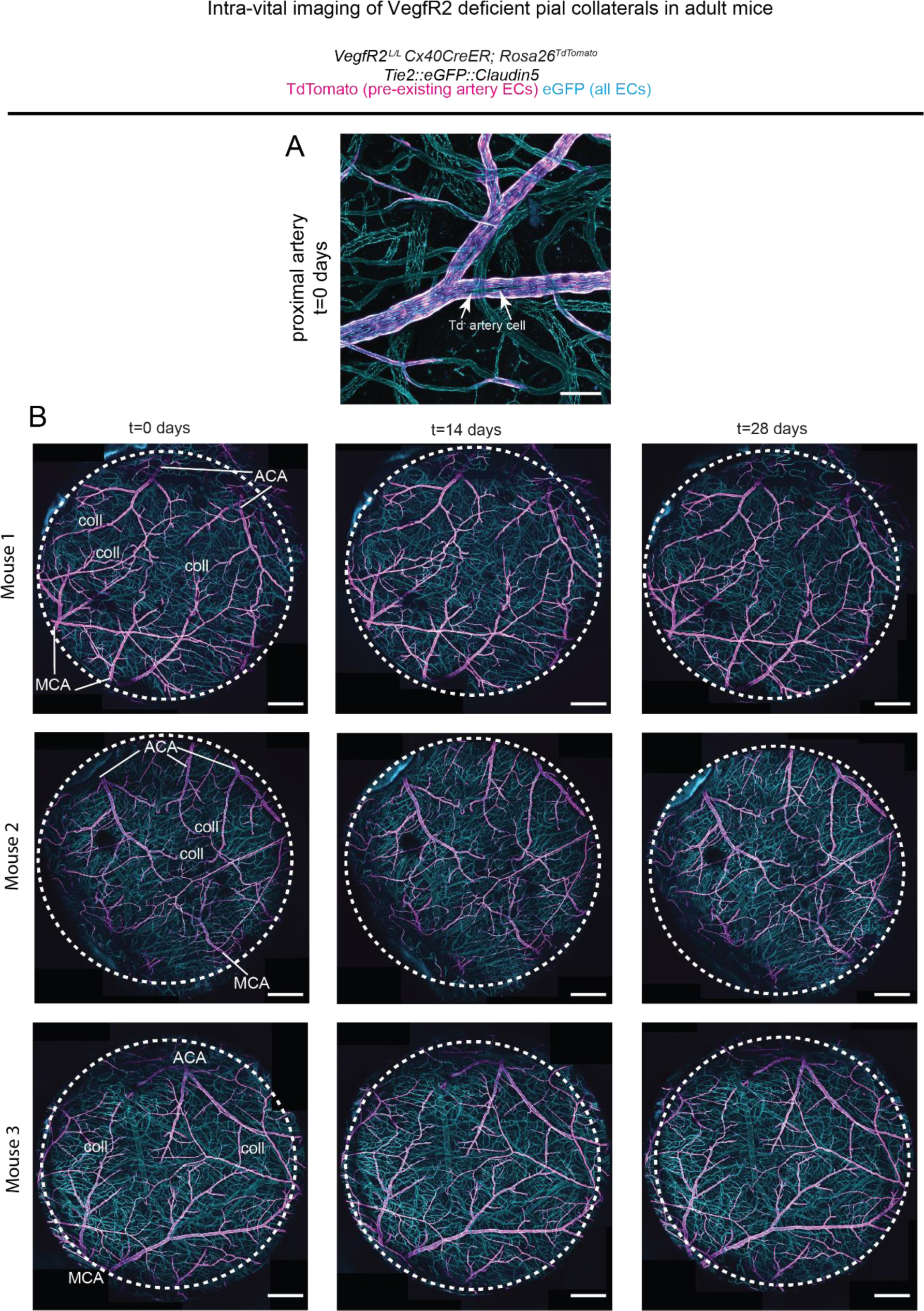
VegfR2-independent maintenance of pial collaterals, related to Figure 6. (**A**) Confocal image of a proximal artery from VegfR2 artery specific knockout adult mouse shown in Figure 6. Arrows point to TdTomato^-^ cell. (**B**) Confocal images of the entire cranial window from 3 different mice show that the overall pial artery network remains unchanged without arterial *VegfR2*. MCA, middle cerebral arteries, ACA, anterior cerebral arteries, Coll, collateral arteries between MCA and ACA. Scale bars: **A**, 100µm; **B**, 500µm.

